# Connectome of a human foveal retina

**DOI:** 10.1101/2025.04.05.647403

**Authors:** Yeon Jin Kim, Orin Packer, Thomas Macrina, Andreas Pollreisz, Christine A. Curcio, Kisuk Lee, Nico Kemnitz, Dodam Ih, Tri Nguyen, Ran Lu, Sergiy Popovych, Akhilesh Halageri, J. Alexander Bae, Joe Strout, Stephan Gerhard, Robert G. Smith, Paul R. Martin, Ulrike Grünert, Dennis M. Dacey

## Abstract

The fovea is a unique specialization of the primate retina and is a promising site for obtaining the first complete connectome of a human central nervous system (CNS) structure. Within the fovea, neural cells and circuits have been miniaturized and compressed during evolution to sample the visual image at highest spatial resolution and begin the neural processing that serves human form, color, and motion perception. Here we present a comprehensive analysis of a sample of human foveal retina using deep learning-based segmentation to reconstruct all cells and synaptic connections at nanoscale resolution. We classified ∼3,000 cells into 51 distinct morphological types based on their structural features and connectivity patterns. Our observations reveal novel synaptic pathways absent in non-human primates, suggesting specialized circuits contribute uniquely to human trichromatic color vision. A biophysical model of the distinct connectomes made by gap junctions (electrical synapses) between short- (S) and medium-long- (ML) wavelength-sensitive cone photoreceptors, suggests chromatic interactions between S and ML cones prior to the first chemical synapse. Segmentation of retinal ganglion cells (RGCs) suggests the presence of only 11 visual pathways, with 5 high-density RGC pathways accounting for over 95% of foveal output to the brain: a dramatic contrast to the 40+ ganglion cell types recognized in mouse retina. Our connectomic analysis reveals distinctive features of human neural circuitry and demonstrates how AI-based computational approaches can advance understanding of human brain structure and function.

**Significant statement:** Vision begins in the retina, an accessible outpost of the brain located at the back of the eye. When we directly view an object, we use the fovea—a tiny region containing over 50 distinct, densely packed neuron types. How these diverse cells interconnect through synapses to transmit visual information has been difficult to determine. Here, we used artificial intelligence-based computational methods to create a complete wiring diagram—a connectome—mapping every neuron and synapse in a sample of human fovea. This connectomic resource will be freely available to researchers worldwide. Discoveries so far include synaptic circuits linked to human color vision, demonstrating how connectomics can reveal what makes the human nervous system unique.

## Introduction

The vertebrate retina is an accessible central nervous system (CNS) outpost where the visual process begins. It has long been appreciated for its great complexity(1) and for linking neural structure to visual function(2–4). The retina was an early focus of synaptic-level connectomic reconstruction(5–10), and more recently, retinal cell populations have been linked to their molecular profiles(11–14) and the genetics of human disease(13, 15). Yet even for this intensively studied neural system, we lack a full wiring diagram that could explain how vision—one of our most fundamental senses—is initiated.

The recent achievement of a complete connectome of the brain of the fruit fly, *Drosophila melanogaster*(16, 17), and application of connectomic methods to mm-scale volumes of mouse and human neocortex(18–21), have made the connectome of a larger mammalian brain a conceivable goal(18, 19, 22, 23). In the present study, we take advantage of the distinctive neural architecture of the foveal retina to provide a first-draft connectome of this critical locus in the human CNS.

The fovea is a specialized central region in the retina of humans and other diurnal primates where neural circuits are spatially compressed to facilitate high-resolution form, color and motion perception. At barely 1 mm wide, the fovea encodes only a few degrees of visual space, but its representation is magnified 100-fold across the surface of primary visual cortex(24–28). Accordingly, the fovea is essential for normal vision, and failure of foveal function is a hallmark of devastating blinding retinal diseases. Understanding the fundamental circuit architecture underlying foveal function therefore represents a critical goal.

From a signal-processing viewpoint, the fovea is an input-to-output sensory structure, where light transduction by photoreceptors is processed by diverse neural cell types across two narrow cell and synaptic layers. Here we provide a catalog of human foveal cells and their synaptic assignments to distinct functional circuits. We use this dataset to directly address the long-standing question of the number and identity of visual pathways that link the fovea to the brain. We define only 11 visual pathways, with five accounting for over 95% of foveal output to the human brain, a striking reduction compared to non-primate mammals(29–31) and non-human primates(32, 33). In addition, we observed unexpected differences in synaptic connections compared to non-human primates and provide evidence that neural coding for trichromatic color vision begins with circuitry that is uniquely human.

## Results

### Acquisition of the foveal volume

Our goal was to obtain complete retinal circuitry near the peak of cone density (peak of visual acuity)(34, 35). This is a difficult task because foveal cones feed inner retinal circuits via very long axons (Henle fibers, Fig. 1A-1C)(36). Our data sample was therefore centered at 500 µm retinal eccentricity, where cone axon length averages 300-400 µm(37, 38); this location corresponds to cones located ∼100-200 µm from the center of the foveal pit (∼0.5-1 deg). We cut 3028 vertical sections at 50 nm thickness through a depth of ∼150 µm and imaged an area 180 µm tall x 180 µm wide at 5 nm resolution (voxel size, 5 x 5 x 50 nm^3^), extending from the Henle fiber layer (outer retina) to the nerve fiber layer (inner retina) (Fig. 1D). Deep learning-based methods (convolutional neural networks)(39, 40) were used to align the images, detect membrane boundaries, segment neurons, glia, blood vessels, conventional (inhibitory) and ribbon (excitatory) synapses, and predict synaptic relationships in the outer and inner synaptic layers (Fig. 1D-1K; see Methods for procedures). We named this volume HFseg1 (Human Fovea, segmented 1). All annotated cell types and the draft catalog of synaptic connections are openly available for exploration using search tools at https://neuromaps.app/datasets (see *SI Appendix*, Table 3), which utilizes the Neuroglancer browser and CAVE infrastructure. In addition, we created a second site, accessed via NeuroMaps, that facilitates proofreading of cells and synapses by the user community.

**Fig. 1.**
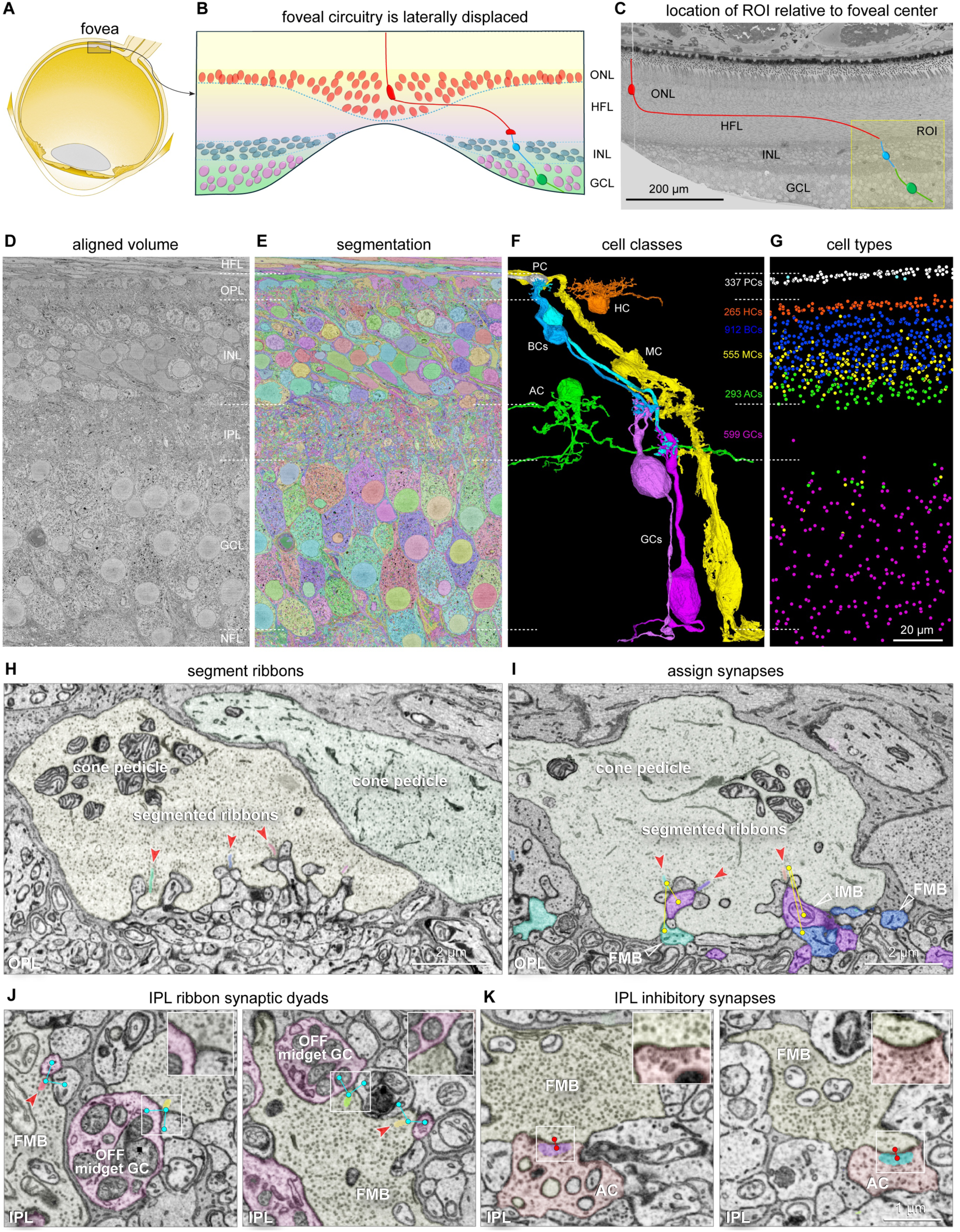
The HFseg1 volume. **A.** The fovea forms a pit in the retina ∼3 mm temporal to the optic disc (within the inset rectangle). **B**. Schematic foveal pit showing cone (red) to bipolar cell (blue) to ganglion cell (green) circuit. ONL (outer nuclear layer), HFL (Henle fiber layer), INL (inner nuclear layer), GCL (ganglion cell layer). **C**. Trimmed retinal block face and ROI (yellow box) in temporal retina 400-500 µm from the foveal center. **D**. Single section from the imaged volume (cropped laterally) to illustrate the retinal layers. HFL, OPL (outer plexiform layer), INL, IPL (inner plexiform layer), GCL, and NFL (nerve fiber layer). **E**. Volume segmented into membrane-bound objects. **F**. Examples of segmented cells: photoreceptor axon terminal (PC, white), horizontal cell (HC, orange), ON and OFF bipolar cells (BC, light and dark blue), Müller glia cell (MC, yellow), amacrine cells (AC, green), ON and OFF ganglion cells (GC, dark and light violet); astrocytes and microglia not shown here. **G**. Numbers and soma locations of the cell types shown in **F**. **H.** Three segmented ribbons (varied colors) at synaptic triads (red arrowheads) in a single cone pedicle. **I.** Synaptic assignments associated with each ribbon (yellow dots and lines). An IMB (inner-ON midget bipolar, violet, white open arrowhead) cell makes an invaginating contact and two FMB cells (outer-OFF midget bipolar, light and dark blue, white open arrowheads) make a basal contact with the cone pedicle. **J.** Examples of ribbon and synapse prediction in the inner plexiform layer (IPL) for an FMB cell axon terminal at two locations (left and right panels). Segmented ribbons (varied colors, red arrowheads) and synaptic predictions (blue dots, lines) are shown. **K.** Inhibitory vesicle cloud segmentation (varied colors) and synaptic prediction (red dots and lines) between an amacrine cell and an FMB axon terminal in the IPL. Insets in **J** and **K** show zoomed view of synapses indicated by white boxes.

### Cell populations

To derive and understand any connectome, the identity and layout of all cell types must first be clearly understood. Here, we took advantage of the detailed anatomical picture of foveal cell types(41, 42) that has been linked to transcriptomically defined cell clusters in human and non-human primate retina(12, 13, 43). Moreover, for many cell populations the presence of orderly spatial arrays could be used to estimate relative density. We were therefore able to use the original segmentation followed by proofreading of cell morphology (∼40,000 edits to date) to annotate and classify all cells and to assess the quality of synapse prediction (Fig. 1F-1G; see *SI Appendix*, Table 1A, 3008 total cells with cell bodies within the volume; ∼304,800 total synapses, including those related to processes extending into or through the volume but not linked to cell bodies).

In brief, HFseg1 contained 313 cone (pedicles) and 24 rod (spherules) synaptic terminals (rod: cone = 0.07, *SI Appendix*, Table 1C, 1D). 17 cone pedicles were identified as short wavelength-sensitive (S) cones (Fig. 2A)(44). The remaining 296 cones comprised long (L) and medium (M) wavelength-sensitive cones, here referred to collectively as LM cones. Postsynaptic to cones, we annotated 268 horizontal cells and 912 bipolar cells (*SI Appendix*, Table 1E, 1F). It was possible to distinguish the H1 and H2 horizontal cell types (Fig. 3) and most of the classically recognized bipolar cell types (Fig. 4) by distinctive morphology and connectivity. Some exceptions will be considered further below.

**Fig. 2.**
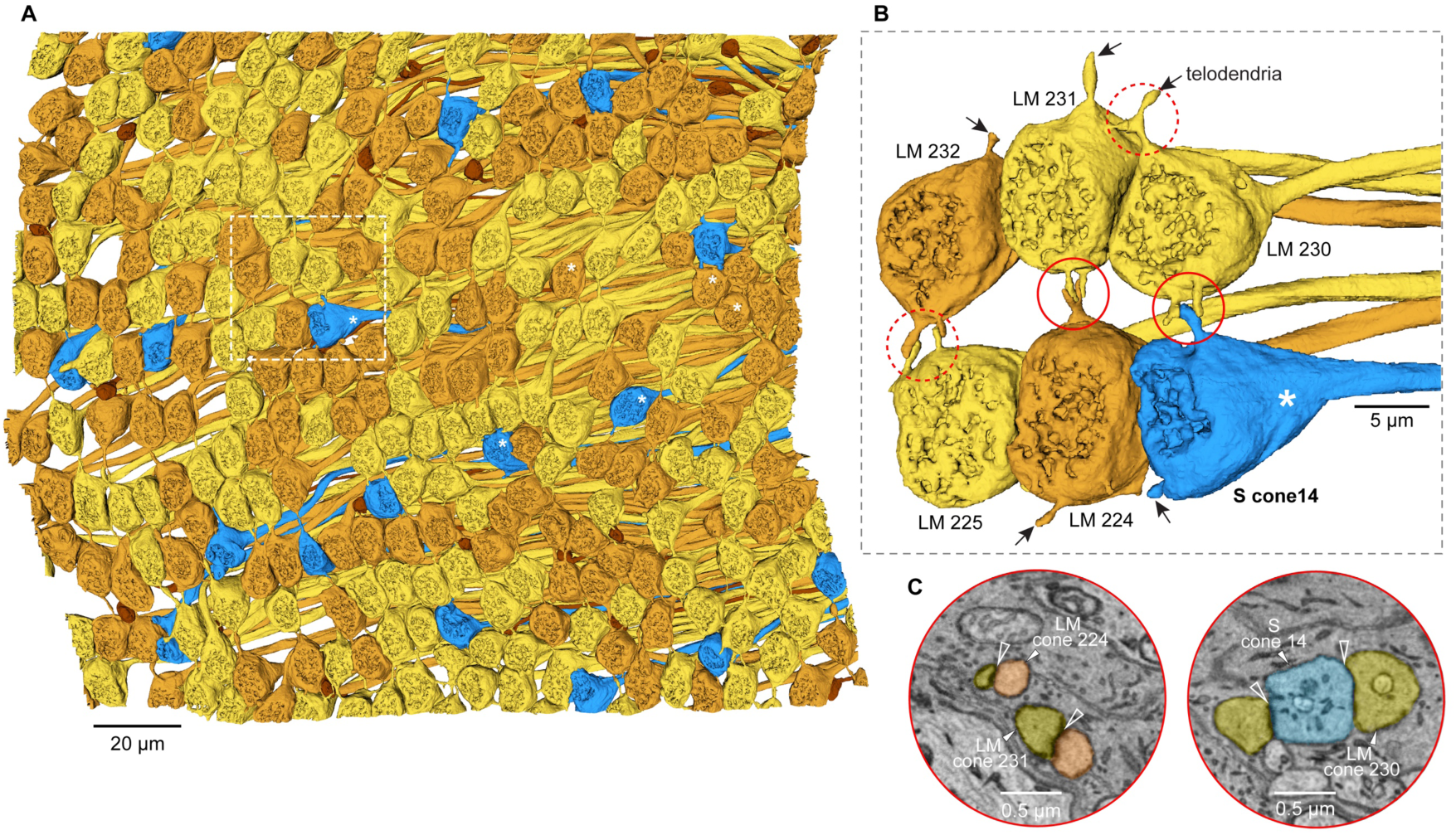
Photoreceptors. **A**. Array of photoreceptor axon terminals (cone pedicles, rod spherules) viewed from the synaptic face of the terminals (296 LM cones, gold; 17 S cones, blue; 24 rods, brown) (see *SI Appendix*, Table 1A, 1C, 1D). Five cones marked with white asterisks were used to generate the vertical connectome analysis shown in *SI Appendix*, Figs. S9 and S10 (S cones 54 and 103; LM cones 21, 22, and 58). **B**. Six cone pedicles within the white dotted box in **A**. These pedicles contact each other via telodendria (e.g., arrows) within the circled areas. One pedicle, identified as S cone 14 (white asterisk), contacts neighboring LM cone 230. **C**. EM micrographs showing telodendritic contacts (open white arrowheads) corresponding to the two solid red circled areas in **B**. **Left**: contact between LM cones 231 and 224. **Right**: S cone 14 contact with LM cone 230. Cone-cone contact surface areas were calculated and used in the simulation shown in Fig. 8.

**Fig. 3.**
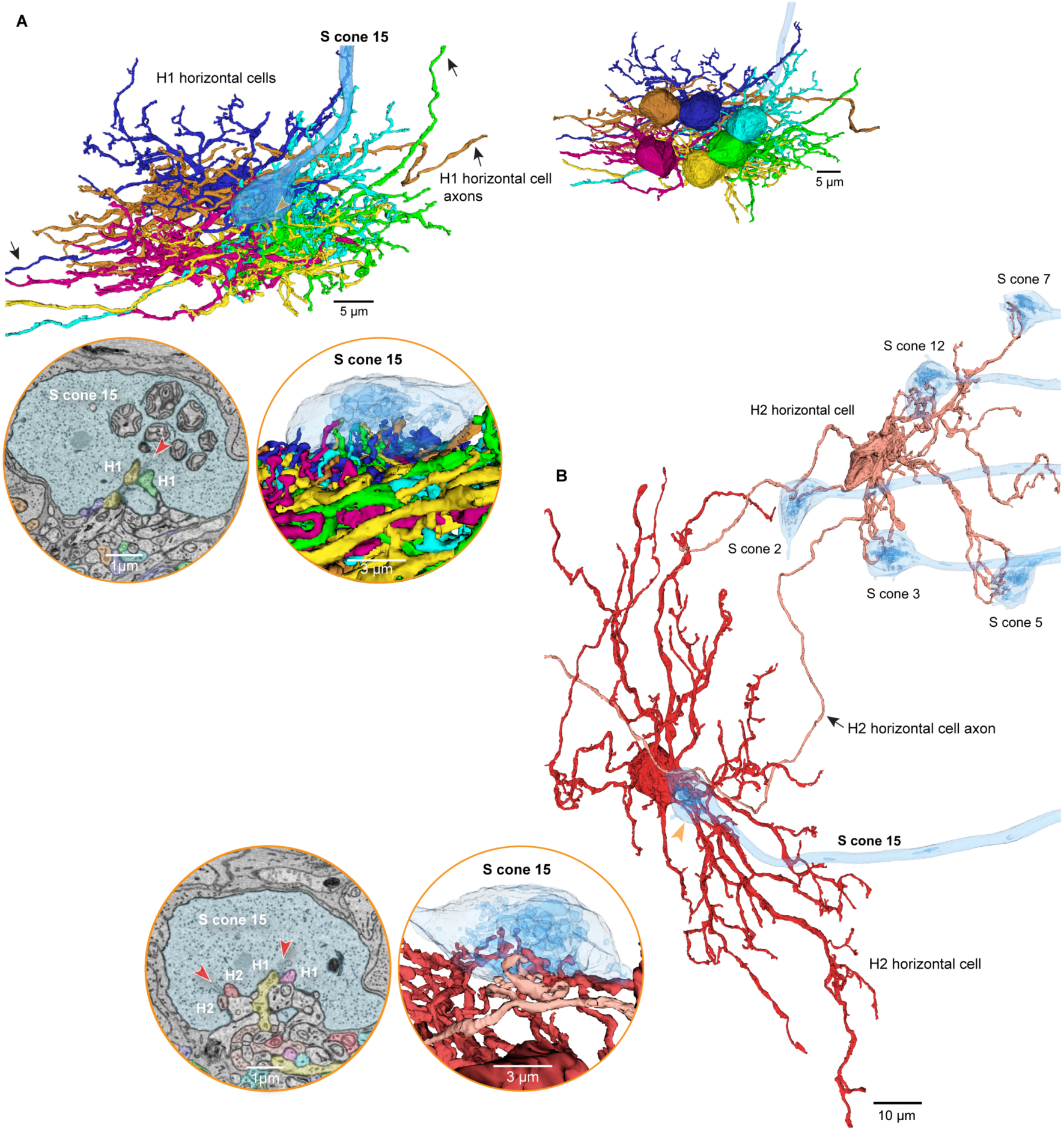
H1 and H2 horizontal cells synapse with S cones. **A.** Six overlapping H1 cells are shown (varied colors); each H1 cell forms lateral elements at S cone 15 (blue). Upper right inset shows an inverted view of the cell bodies of the six H1 cells. Lower left inset shows EM view of H1 cell lateral elements at a synaptic invagination of S cone 15 (red arrowhead). Lower right inset shows a 3D view of S cone 15 (partial transparency) contacting multiple H1 cells. H1 cells made 43% ± 16% of the total contacts with S cones (n = 10 S cones). **B**. Two H2 cells (tan and red) contact multiple S cones, including S cone 15. Lower insets show H2 cell lateral elements at S cone 15 in single-layer EM (left) and 3D (right) views (as in **A**). H2 cells made 57% ± 16% of the total contacts with S cones (n = 10 S cones).

**Fig. 4.**
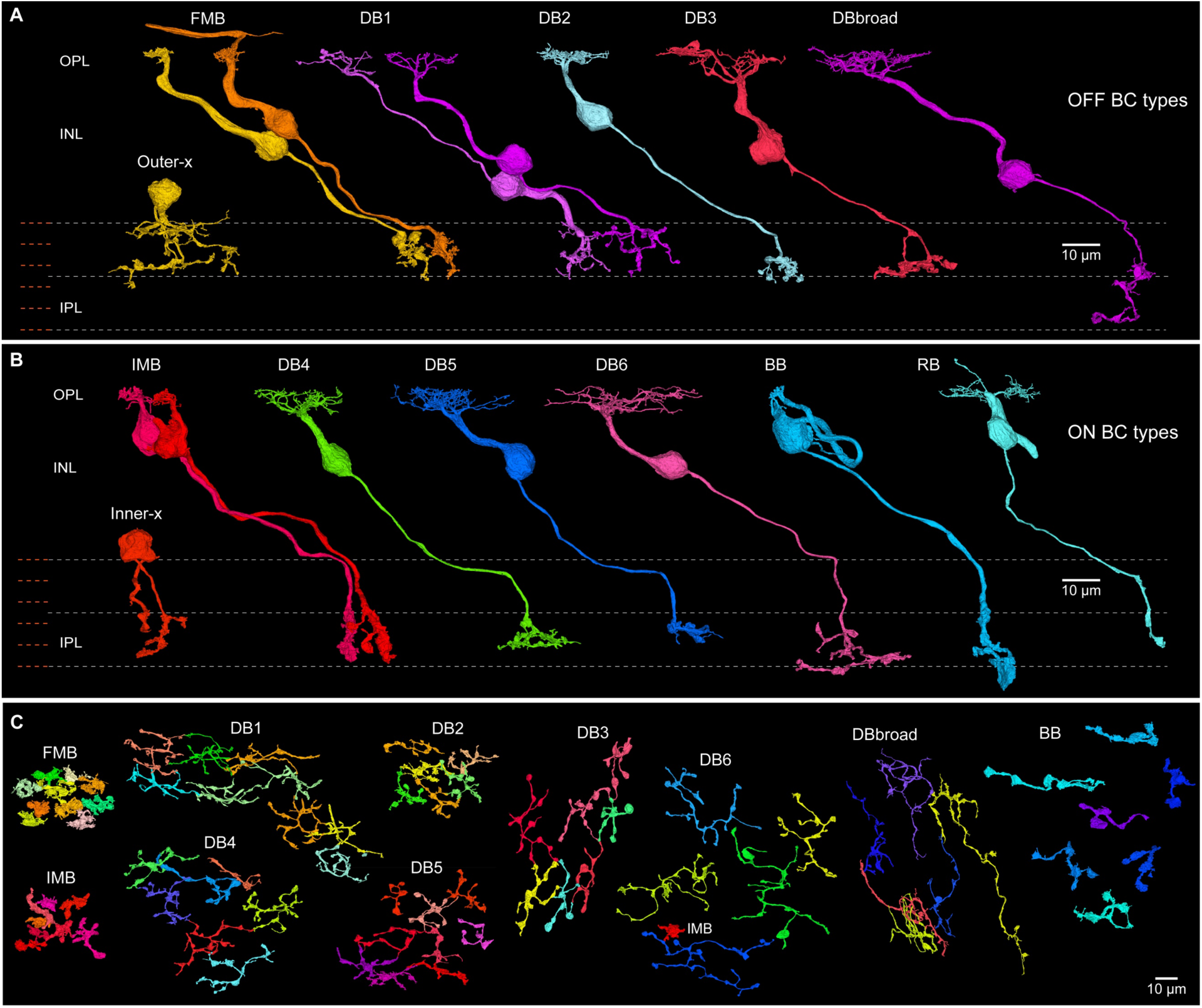
Bipolar cell types. **A**. Outer stratifying types: FMB (flat midget bipolar), DB (diffuse bipolar types). Outer-x bipolar cells lack dendrites but make ribbon synapses in the IPL (*SI Appendix*, Fig. S6A). DBbroad cells extend processes across the IPL depth (*SI Appendix*, Fig. S7). **B.** Inner stratifying bipolar types: IMB (invaginating midget bipolar), DB (diffuse bipolar types), BB (blue-cone bipolar), RB (rod bipolar). Inner-x bipolar cells are shown in *SI Appendix*, Fig. S6C. **C**. Horizontal view of spatial mosaics of axon terminals for small groups of neighboring cells of each bipolar type, as labeled.

We annotated 293 amacrine cells, including 40 cells displaced to the ganglion cell layer (Fig. 5; *SI Appendix*, Table 1G; *SI Appendix*, Fig. S8), first dividing cells by dendritic field diameter into small (diameter < 100 µm; n = 189), and large-field (diameter > 100 µm) groups (n = 104). We divided these groups in turn into 19 types (6 small-field and 13 large-field; *SI Appendix*, Table 1G) based on morphology, stratification, mosaic interactions and nuclear morphology (*SI Appendix*, Figs. S3, S6, S8). Figure 5 provides a partial view of this dataset by highlighting four cell types (see also *SI Appendix*, Table 3); two additional amacrine populations are illustrated in *SI Appendix*, Fig. S8 and others can be viewed at the NeuroMaps site.

**Fig. 5.**
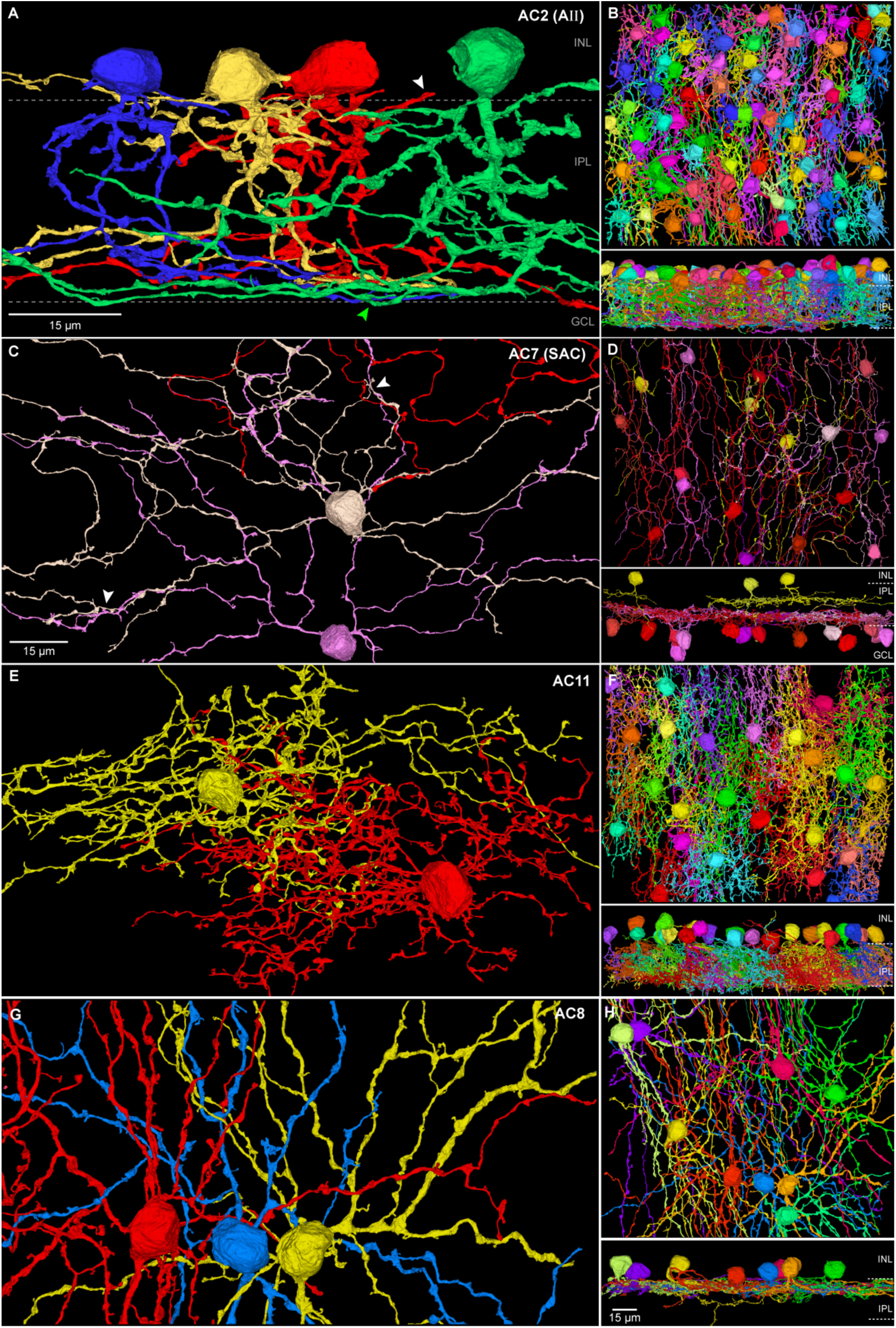
Amacrine cells. Amacrine cells in the INL (253 cells) and GCL (40 cells) were divided into 19 types (AC1-19; *SI Appendix*, Table 1G). **A.** Small-field Aλλ amacrine cells (AC2) shown in vertical view, with outer lobular dendrites (white arrowhead) and extensive inner dendrites (green arrowhead). **B.** Horizontal view of the entire Aλλ population in HFseg1. Lower panel: vertical view. **C.** Starburst amacrine cells (AC7) were located primarily in the GCL (14 cells; INL, 3 cells). White arrowheads indicate typical fasciculation of starburst dendrites. **D.** Mosaic of starburst cells, with GCL cells (red-violet) and INL cells (yellow); lower panel: vertical view. **E**. Small-field amacrine cell type (AC11) in horizontal view with densely branched dendritic arbor. **F**. Horizontal view of the AC11 cell mosaic; lower panel: vertical view showing dendrites that extend broadly across the IPL. **G.** Large-field amacrine cells (AC8) with thick lobular dendrites, narrowly stratified in the outer IPL. **H.** Horizontal view of the complete mosaic; lower panel shows vertical view of outer IPL stratification. Scale bar in **C** applies to **E** and **G**. Scale bar in lower panel of **H** applies to **B, D**, and **F**.

**Fig. 6.**
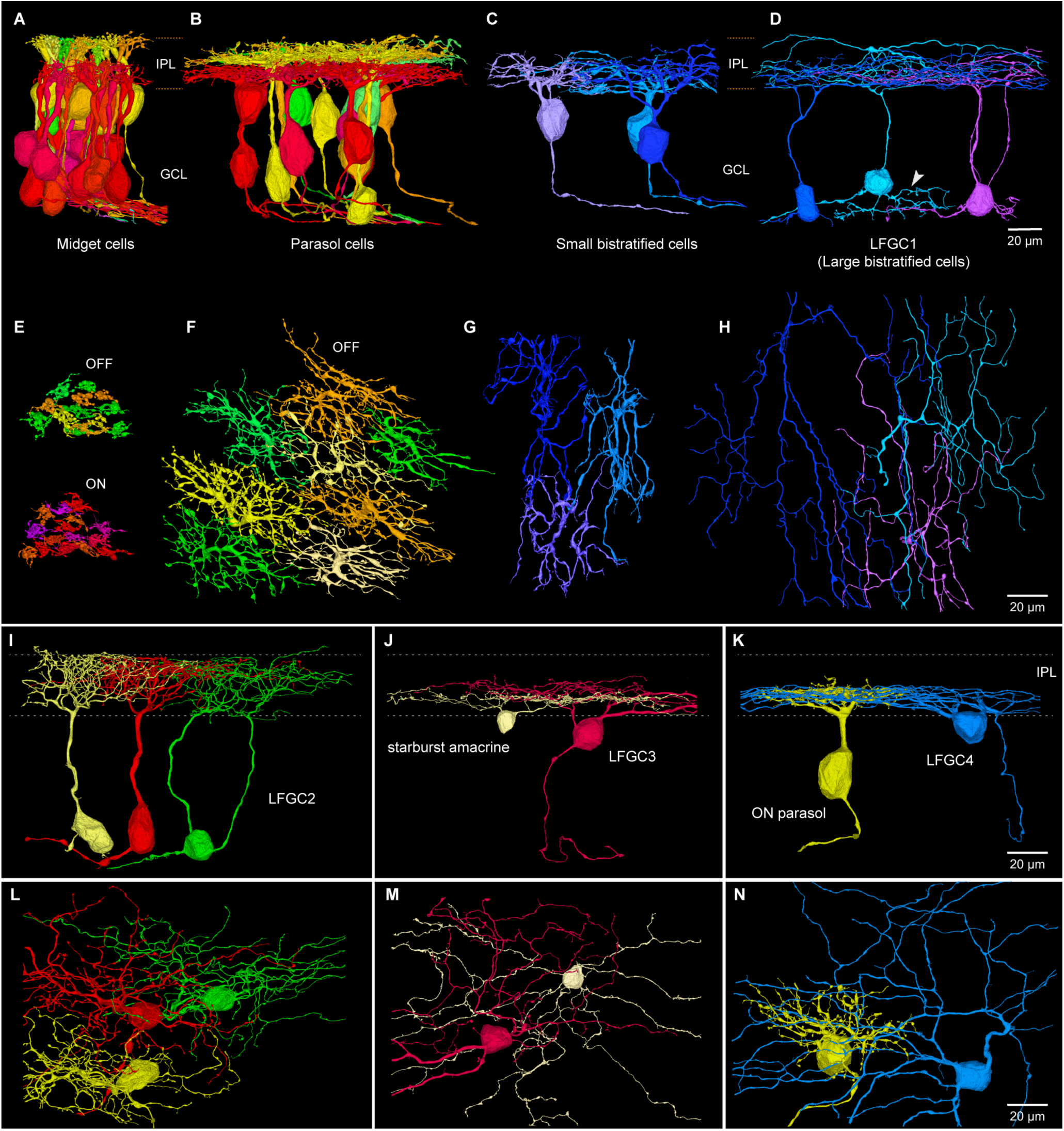
Ganglion cells. **A**. Cluster of inner-ON (red-violet) and outer-OFF (yellow-green) midget ganglion cells; dendrites stratify broadly across inner (ON) and outer (OFF) halves of the IPL. **B**. Inner-ON (red-violet) and outer-OFF (yellow-green) parasol ganglion cells; dendrites stratify broadly near the center of the IPL. **C**. Small bistratified ganglion cells extend dendrites across the ON-OFF subdivision of the IPL. **D**. Large bistratified ganglion cells (LFGC1) show large dendritic fields and broad stratification. White arrowhead shows branching processes of unknown significance arising from the cell body. **E-H**. Mosaics of dendritic arbors in horizontal view of the same cells shown in **A-D**. **E**. Outer-OFF (yellow-green, top) and inner-ON (red-violet) midget GCs. **F**. Outer-OFF parasol cell dendrites show minimal overlap (inner-ON parasol mosaic not shown). **G**. Small bistratified cells show minimal dendritic tree overlap. **H**. LFGC1 (Large bistratified cells) show a much larger and more sparsely branching irregular tree. **I-K**. Morphology of three additional large field types. **I**. LFGC2 (Large diffuse cells) show a densely branched tree broadly stratified across the IPL. **J**. LFGC3 (Recursive bistratified cells) are narrowly bistratified near the center of the IPL and tend to fasciculate with starburst amacrine cells. **K**. LFGC4 (Inner smooth monostratified cells) show large, monostratified dendritic trees costratified with parasol cells. **L-N**. Horizontal views of the cells shown in **I-K.**

We annotated 599 ganglion cells (Fig. 6; *SI Appendix*, Table 1H). As expected, the classically recognized midget GCs constitute the great majority (536 cells, ∼90%)(13, 45); these cells were readily divided into outer-OFF stratifying (n = 280 cells) and inner-ON stratifying (n = 256) types (Fig. 6A, 6E). A second well-defined cell group, the parasol cells (26 cells, 13 inner-ON cells and 13 outer-OFF cells, ∼4% of total GCs), was also easily distinguished (Fig. 6B, 6F). Small bistratified GCs(46), found to show blue-yellow color opponency in macaque retina(47, 48) were distinguished by their broad dendritic ramification (12 cells, ∼2% of total GCs; Fig. 6C, 6G) and extensive synaptic input from blue cone bipolar cells (BB cells)(49). Together, the midget, parasol and small bistratified cells accounted for 95.8% of the ganglion cells in HFseg1 (*SI Appendix*, Table 1H). The remaining 25 ganglion cells showed relatively large dendritic fields (Figs. 6D, 6H–6N) and were divided into 5 provisional groups comprising 6 types (*SI Appendix*, Table 1H; Supporting Information Text) pending a more detailed analysis of their synaptic connections, giving a total of 11 ganglion cell populations in this volume–this number is very close to the 12 populations identified for central human retina by molecular profiling(13).

Finally, glial cells were readily distinguished from all neurons by their distinctive ultrastructure (*SI Appendix*, Figs. S3A, S3B, S3C, S3F). The Müller cells (555 cells, Fig.1F, 1G; *SI Appendix*, Table 1B) were distinguished as radial glia with cell bodies in the inner nuclear layer (*SI Appendix*, Fig. S1A). A second, morphologically distinct population of glial cells in the GCL was closely associated with superficial blood vessels (*SI Appendix*, Figs. S1B, S3B) and identified as astrocytes(50) (32 cells, *SI Appendix*, Table 1B). We also identified 15 microglial cells (*SI Appendix*, Table 1B) with cell bodies in the IPL or GCL (*SI Appendix*, Figs. S1A, S3A). The density of microglial cells was low, consistent with previous measurements in the macaque monkey foveal retina(51, 52). In the current volume, we have not yet attempted to characterize blood vessel and associated pericytes, though these cellular elements are also segmented and are available to annotate and further characterize. More detailed information on cell type identification is provided in the supporting information (Supporting Information Text on Cell type identification).

### Synapse detection

To assess the accuracy of ribbon segmentation and synapse prediction (Fig. 1H-1K), we compared deep-learning-predicted synaptic assignments to manual (ground truth) assignments. We focused on the cone-to-midget bipolar cell synapse in the OPL and the midget bipolar cell synapses to amacrine and ganglion cells in the IPL. For ground truth, we manually annotated all midget pathway synaptic connections for three LM cones (cones 21, 22, and 58; *SI Appendix*, Fig. S9; *SI Appendix*, Table 2A-2C).

Figure 7 shows the predicted outgoing synapses for a pair of midget bipolar cell axon terminals. We calculated precision (true synapses / [true synapses + false positives]; the accuracy of positive predictions) and sensitivity (true synapses / [true synapses + false negatives]; the proportion of true synapses identified) for ribbons and their synapses. The segmentation of ribbons showed high precision and sensitivity near 100% (precision: 93.5% ± 2.7%; sensitivity: 99.7% ± 0.8%, n = 6 midget bipolar cells, 370 total ribbons; Fig. 7B inset). Postsynaptic partner assignment for these ribbons was also excellent (precision: 89.0% ± 5.4%; sensitivity: 93.7% ± 4.6%, n = 6 midget bipolar cells, 637 total synapses; Fig. 7B inset). These values demonstrate the high accuracy of convolutional nets for predicting and segmenting ribbon synapses.

**Fig. 7.**
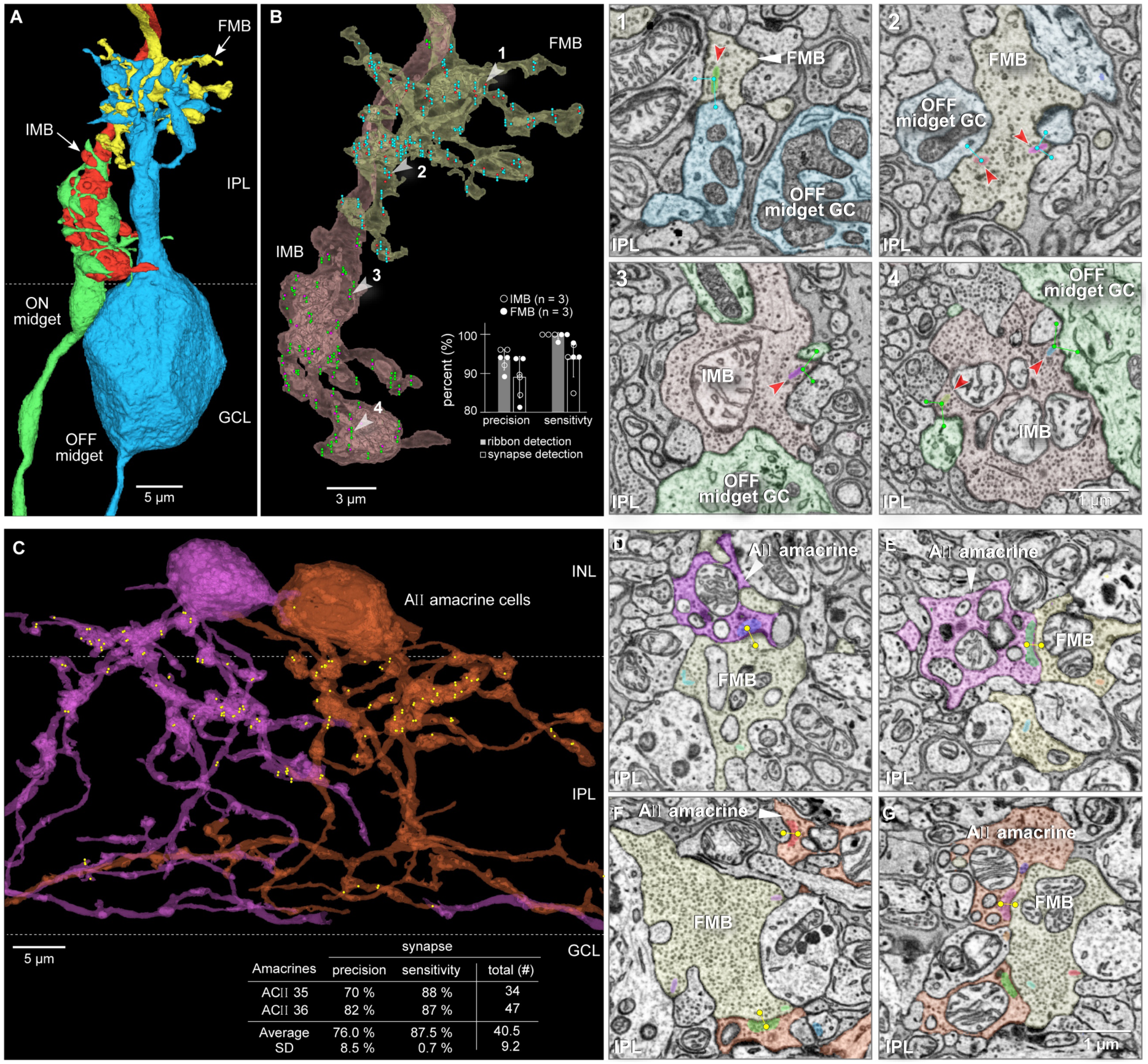
Synapse detection. **A**. OFF midget bipolar (yellow) and ON midget bipolar (red) presynaptic to an outer-OFF (blue) and inner-ON (green) midget ganglion cell. **B.** Outgoing synapses detected for the midget bipolar cell axon terminals shown in **A** (ganglion cells removed, semi-transparent view). Synaptic ribbons (red/violet dots) and postsynaptic profiles were first identified manually for cone 22, BC 308 (IMB), and BC 303 (FMB) (see *SI Appendix*, Fig. S9). Predicted outgoing synapses are shown as green dots and lines (IMB) or blue dots and lines (FMB). White arrowheads numbered 1-4 indicate the panels shown on the right in single-layer EM view, displaying segmented synaptic ribbons (in varied colors, red arrowheads) and predicted postsynaptic segments (dots and lines linking ribbons to postsynaptic structures); unlabeled postsynaptic profiles belong to amacrine cells. Inset histogram shows precision and sensitivity of ribbon detection and synaptic assignment. **C.** Inhibitory output synapses made by two neighboring Aλλ amacrine cells (purple and orange, shown in partial transparency). Yellow dots show predicted synaptic outputs; most synaptic outputs arise in the outer half of the IPL and are presynaptic to OFF bipolar cells. Inset shows precision and sensitivity for the synapses made by each Aλλ cell. **D-E**. Single-layer images showing segmented vesicle clouds (varied colors) in the purple Aλλ cell (shown in **C**) and predicted synaptic connections with an FMB cell (yellow dots and lines); segmented ribbons in the FMB are also shown. **F-G**. Single-layer images showing segmented vesicle clouds (varied colors) in the orange Aλλ cell (shown in **C**) and predicted synaptic connections with another FMB cell (yellow dots and lines); segmented ribbons in the FMB are also shown.

**Fig. 8.**
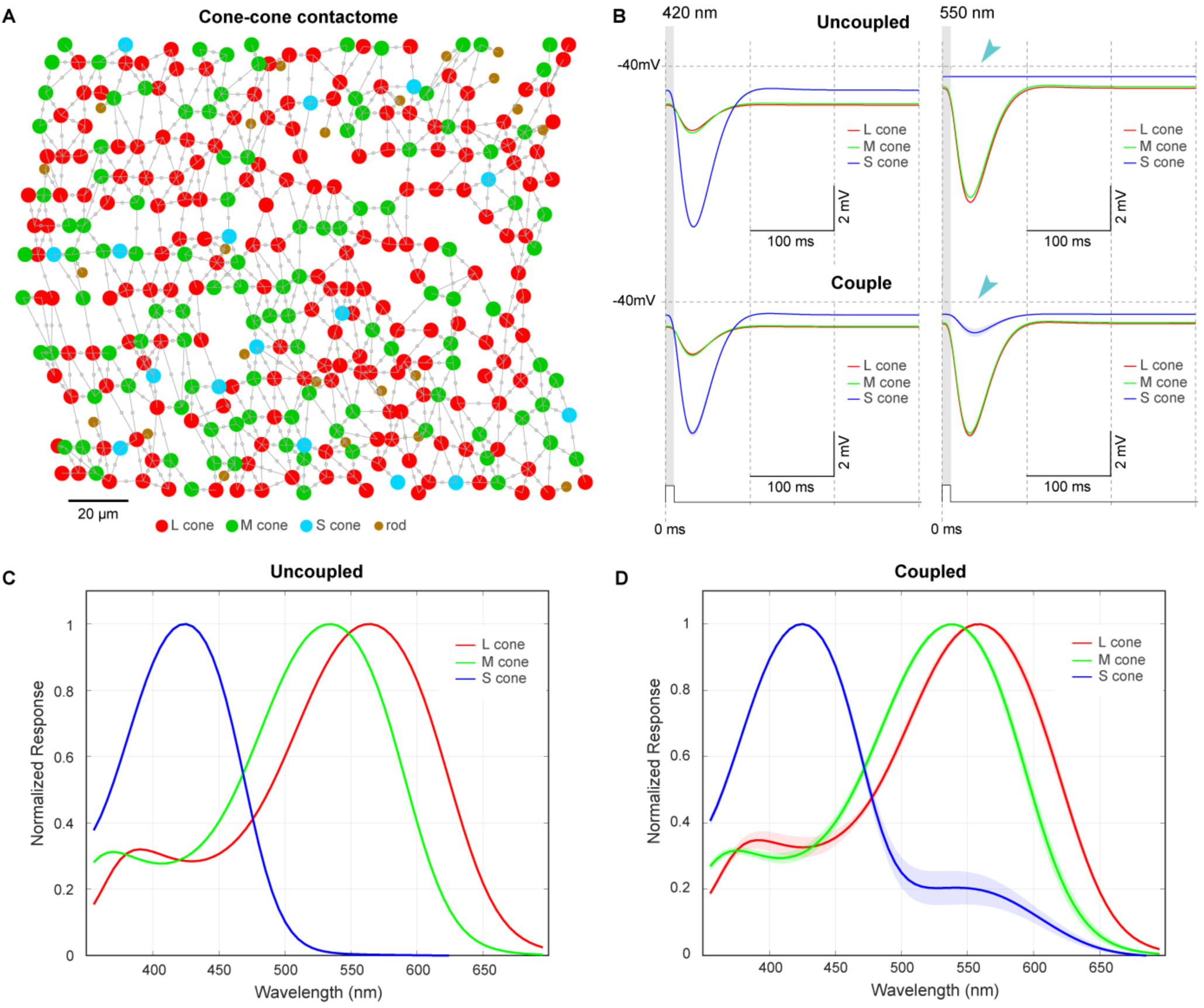
Cone-cone coupling alters S cone spectral tuning. **A**. Biophysical model of the cone-cone connectome. Assignment of L/M cone type was random with a ∼1.7:1 L:M ratio, based on known proportions in the human fovea. Contacts between cones were determined from the reconstructed connectome, and conductances were assigned based on the contact area, known gap junction density, and single connexin channel conductance. Cones were modeled compartmentally as a soma that included the phototransduction cascade, connected to an axon, in turn connected to the axon terminal (see model parameters in *SI Appendix*, Table 4). **B**. Modeled voltage responses (averages from 5 different L, M, and S cones in the network) to monochromatic 10 msec light flashes at 420 nm and 550 nm. **Top (Uncoupled):** zero cone-cone conductance. **Bottom (Coupled)**: cones coupled via individual cone-cone conductances given in the electrical model. Coupling among L, M, and S cones produces a light-evoked response in S cones at 550 nm (teal arrowhead) that is absent in the uncoupled condition. **C**, **D**. Impact of cone-cone coupling on the action spectrum of the cone voltage response. Average light responses were generated every 5 nm from 350 to 700 nm. **C** (Uncoupled condition): the action spectrum is proportional to the spectral sensitivity of the L, M, and S cone opsins. **D** (Coupled condition): the S cone voltage response shows a smaller second peak of sensitivity at ∼550 nm (teal arrowhead), which is not present when cones are uncoupled.

In contrast to the ribbon synapses made in the IPL, ribbon synapses made by the cone pedicle in the OPL are extremely complex(53, 54) (Fig. 1H, 1I). Postsynaptic processes can either invaginate into the face of the pedicle or occupy a basal position at the cone pedicle up to ∼800 nm from the sites of transmitter release(54, 55). We mapped all cone-to-bipolar synapses at the dendritic tips for a sample of six midget bipolar cells. We found high precision and sensitivity for ribbon detection (n = 51 ribbons) (precision: 92.3% ± 6.7%; sensitivity: 100%, n = 3 LM cones, *SI Appendix*, Fig. S2A-S2C) and for the postsynaptic processes (n = 153) deriving from the invaginating (IMB) and flat (FMB) midget bipolar cells (precision: 98.7% ± 1.2%; sensitivity: 99.7% ± 0.6%, n = 153 total synaptic connections; see *SI Appendix*, Fig. S2C right). Diffuse bipolar (DB) cells, which make sparser connections with multiple cones, were also sampled (n = 2, DB4 and DB2; *SI Appendix*, Fig. S2D-S2F), yielding similar precision and sensitivity values.

For inhibitory synapses, we focused on the Aλλ amacrine cell (Figs. 5A, 7C, 9; *SI Appendix*, Table 1G) where glycinergic synapses to OFF bipolar cell axon terminals are well established(56–59). We manually mapped all the outgoing synapses made by two neighboring Aλλ amacrine cells and compared our ground truth data to the predicted synaptic assignments (Fig. 7C-7G). The prediction captured well the synaptic output from the outer stratifying lobular appendages of the Aλλ cell to outer-OFF bipolar cells (Fig. 7D-7G). Vesicle clouds were detected with high sensitivity (87.5% ± 0.7%, n = 2 Aλλ cells; false negatives were rare; Fig. 7C inset), but relative to ribbon detection, there were more false positives, thereby reducing precision (76.0% ± 8.5%, n = 2 Aλλ cells). Despite this limitation, the results show that our dataset predicts synaptic connectivity with good sensitivity and precision. However, we implemented methods within the Neuromaps interface to proofread all synapses for accuracy.

**Fig. 9.**
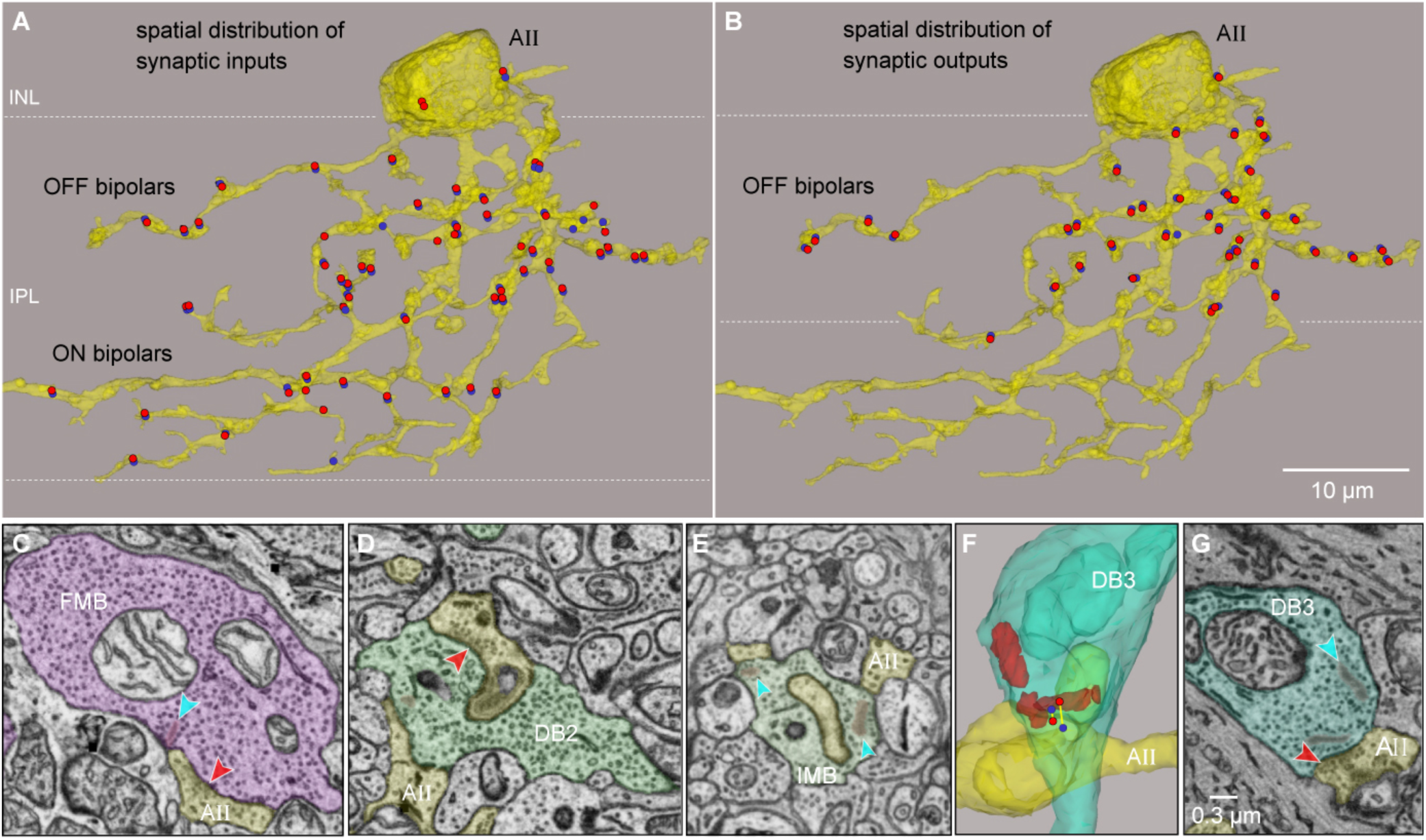
**A**λλ **cells are postsynaptic to both ON and OFF cone bipolar cells at ribbon synapses. A**. Chemical (ribbon) synaptic inputs from multiple OFF and ON bipolar types. For this cell, 70 synapses (red dots, presynaptic locus; blue, postsynaptic locus; most blue postsynaptic dots are overlying the presynaptic position and are not visible in this zoomed-out view) are dispersed throughout the dendritic tree. Across 6 Aλλ cells analyzed, 300 synapses arose from all ON and OFF cone bipolar types, with IMB and FMB cells accounting for 19% and 24% of the synapses, respectively. **B**. By contrast, this Aλλ cell is presynaptic only to OFF cone bipolar cells; many of these synapses are reciprocal. **C**. An OFF-midget bipolar (FMB, violet) makes a ribbon dyad synapse (blue arrowhead points to ribbon) with the Aλλ cell (yellow) and a parasol ganglion cell. The Aλλ cell makes a nearby reciprocal synapse (red arrowhead) onto this FMB. **D.** Nonreciprocal Aλλ synapse with a DB2 OFF bipolar type (teal). **E.** Aλλ cell (yellow) postsynaptic to an ON-midget bipolar (IMB, green) at two dyad ribbon synapses (blue arrowheads). **F**. 3D view of a reciprocal synapse between this Aλλ cell and a DB3 bipolar (blue); the Aλλ cell inserts a finger-like projection into this bipolar axon terminal. **G**. Single section through contact in **F**, showing two of the three ribbon synapses (blue arrowhead indicates one) and the reciprocal Aλλ inhibitory synapse (red arrowhead).

Having established the accuracy of our connectomic dataset, we examined the cell types and their synaptic connections to identify features that are distinctive in human compared to non-human primates. Below we highlight examples where differences are dramatic and warrant further study.

### Modeling the connectome: S-LM cone electrical coupling alters spectral tuning

Cone pedicles contact each other by short extensions (telodendria) or by distinct pedicle-to-pedicle appositions at perforations in the glial sheath surrounding cone synapses (Fig. 2B). These contacts harbor gap junctions, i.e., electrical synapses that mediate a conductance between cones, including L and M cones(60). In non-human primates, L to M cone contacts are abundant, but S to LM contacts are greatly reduced or absent(44, 61). In the macaque, electrical coupling was absent between LM and S cones(60), consistent with a lack of contacts between these cone types(61, 62). The crosstalk between L and M cones serves to reduce noise in the light-evoked response with a small cost to spectral separation(63–65). The lack of gap junctions between LM and S cones has been interpreted as reflecting the need to keep S and LM cone signals separate to build color-coding receptive fields at later stages in color processing(66, 67).

Unlike in the macaque, S-LM cone contacts have been reported in human retina(44, 68). We therefore mapped the full cone-cone connectome to determine whether S-LM contacts are a consistent feature of the human fovea. Cone-cone contacts (n = 1141) were characterized by a thick, darkly stained region defining the contact area (see Fig. 2C)(69). We found abundant S-LM cone contacts (n = 17 S cones; 5.25 ± 2.11 contacts/S-cone pedicle), indistinguishable from LM-LM cone contacts (n = 296 LM cones; 6.89 ± 3.08 contacts/pedicle). We then used this cone-cone connectome dataset to estimate the effect of S-LM coupling on spectral tuning in a simulated cone-cone network, using known parameters for gap junction number and single channel conductance values as follows. We mapped the surface area of the contacts among all cones in the volume (Fig. 2A), using automated methods to generate the “contactome” between cone pedicles. Detected contacts were inspected and filtered by size to remove very small and likely spurious contacts. The surface area of each contact was calculated (n = 1141 contacts; mean surface area 0.56 ± 0.31 µm^2^). In general, contacts were large and obvious with a relatively thick, darkly stained region defining the contact area (see Fig. 2C), reflecting the presence of large adherens junctions at the contact site(69). We calculated the conductance for each contact based on previously measured densities for gap junctions and the conductance of single gap junction channels (9 gap junctions/µm^2^ at 15 pS/junction; see Methods for details). We used a fixed ratio of L to M cones of ∼1.7:1, randomly assigned(70), and incorporated the cone-cone coupling given by our contactome (Fig. 8A). Each cone was modeled compartmentally as a soma, Henle fiber-axon, and synaptic terminal (see model parameters in *SI Appendix*, Table 4). Our modeled average coupling conductances were low (LM-LM cone pairs, 133 pS, n = 25 cones; S-LM cone pairs, 166 pS, n = 15) by comparison with conductances measured directly between LM cone pairs in the macaque (> 500 pS) or cone pairs in non-primate mammalian retina (∼220-320 pS)(65, 67). Nevertheless, we found that even with this conservative estimate of coupling conductance, the S-cones showed an increased sensitivity to light pulses at longer wavelengths in the model (Fig. 8B). We measured the effect on spectral sensitivity by simulating light responses from L, M, and S cones at 5 nm wavelength intervals across the visible spectrum. The results demonstrate a broad secondary elevation in sensitivity for S cones at longer wavelengths that peaks at ∼550 nm (Fig. 8C-8D) in the coupled cone network. These data provide evidence that S-LM cone coupling can directly influence the spectral tuning of the voltage response of S cones and raise the question of how this network feature affects downstream processing.

### Trichromatic horizontal cells

A second difference in synaptic connectivity that may impact color vision occurs at the synapse between cones and horizontal cells. The horizontal cell network in the outer retina generates lateral negative feedback that underlies the antagonistic receptive field surround of cones, bipolar cells, and ganglion cells(71). In the macaque monkey retina, the horizontal cell-mediated surround is a key element in the circuitry that initiates red-green color vision(72–74). In the peripheral retina of the macaque and marmoset, H1 cells make dense contacts with LM cones but largely or completely avoid contact with S cones, whereas H2 cells show some selectivity for S cones(75–77). While foveal H2 cells in HFseg1 maintained a clear preference for synapsing with S cones (Fig. 3B), we found that H1 cells also received a major input from S cones, comparable to H2 cells (H1, 43% ± 16%; H2, 57% ± 16%, n = 10 S cones). By contrast, H2 cells formed less than 2% of the horizontal cell contacts with LM cones (H1, 98.1% ± 0.1%; H2, 1.9% ± 0.1%, n = 2 LM cones) (see also *SI Appendix*, Fig. S4). Consequently, human foveal H1 cells are predicted to show a trichromatic receptive field with a small S cone contribution (Fig. 3; *SI Appendix*, Fig. S4). Taken together, the S-LM cone coupling and the combined S and LM cone connectivity of H1 horizontal cells provide new connectomic evidence for differences between human and non-human primate circuits that initiate color vision(44).

### Unexpected cell types and circuits in HFseg1

We also found differences in both bipolar and amacrine cell circuits compared to expectations from the macaque monkey, whereby some bipolar cell types were either greatly reduced or absent. For example, DB-OFF bipolar cells can be divided into 4 morphological types (DB 1, 2, 3a and 3b) in macaque monkey(78, 79) but we were only able to distinguish three populations by mosaic arrangement and stratification in the IPL that we call DB1, 2 and 3 (see Supporting Information Text). Bipolar cell morphologies not previously described in primates were also present. One bipolar group showed amacrine-like morphology (outer-x and inner-x, Fig. 4A, 4B) but contained abundant synaptic ribbons (20 cells, *SI Appendix*, Table 1F; *SI Appendix*, Fig. S6). Similar cells were discovered recently in mouse retina(11, 80) and likely correspond to the OFF-x bipolar cells identified by transcriptomics of human fovea(13). While most of these cells stratified in the outer half of the IPL (outer-x cells, n = 15, *SI Appendix*, Fig. S6A-S6B), five cells stratified in the inner half of the IPL (inner-x cells, *SI Appendix*, Fig. S6C-S6G). Another previously undescribed bipolar type showed the characteristic DB cell connection to multiple cones but extended axonal branches in both outer and inner IPL where it provided synaptic output to outer-OFF and inner-ON ganglion cell types (see also *SI Appendix*, Fig. S7C, S7E). We refer to this cell group as DBbroad (18 cells, Fig. 4A; *SI Appendix*, Fig. S7; *SI Appendix*, Table 1F). The HFseg1 volume offers a unique opportunity to further understand these novel bipolar cell populations.

For the amacrine cells, the Aλλ type (AC2) was present at the highest density of any amacrine cell population (26%, 75 cells, Figs. 5A, 5B, 7C, 9; *SI Appendix*, Table 1G). This cell type was recognized by its well-established morphology across species(56), including humans(41, 57, 81). A major function of Aλλ cells is to transfer signals from rod bipolar to cone bipolar cells, serving as conduits for rod signals to ganglion cells at scotopic light levels(82). The presence of Aλλ amacrine cells at such a high density in the fovea, where rods are nearly absent, suggests, however, a functional switch to a primary role in photopic spatial vision. Indeed, we observed that Aλλ cells are postsynaptic to all cone bipolar types, including IMB and FMB cells, across the full extent of the dendritic tree (Fig. 9). These data support the general view that Aλλ cells “have a day job”(57) to provide an inhibitory pathway from ON to OFF bipolar cells under photopic conditions(82–84) while at the same time suggesting specialized synaptic alterations for the Aλλ cells in the human fovea in relation to the midget circuit.

### Visual pathway reduction: the vertical excitatory connectome

Despite its fundamental importance, the number of visual pathways linking the human fovea to the brain is still unknown. Our classification of 11 ganglion cell types suggests that several ganglion cell types observed in the monkey and human retinal periphery are absent from the human fovea. This result also stands in sharp contrast to the retina of the mouse, where ∼40 GC types have been distinguished independent of retinal location(29–31). Our classification, however, did not rule out the possibility that some ganglion cell types with somata outside HFseg1 have dendritic trees which penetrate the volume and contribute substantially to the synaptic connections of the inner retina. We therefore took a second approach to identifying the visual pathways in HFseg1 by mapping what we called the “vertical excitatory connectome”.

We reasoned, as others have(85), that to generate parallel pathways to the brain, each cone photoreceptor must be locally presynaptic to each cone bipolar type (For the S cones there is uncertainty regarding this point, due to their more specialized role in color vision(86, 87)). At the next synaptic step, the combined output of all bipolar cell types should be distributed to all ganglion cell types. In other words, in principle, *all visual pathways should be synaptically linked to the output of any given cone*.

To test this hypothesis, we annotated the ribbons within five cones (three LM and two S) and their postsynaptic bipolar cells. For each bipolar cell, all synaptic ribbons and their postsynaptic GC partners were then identified (Fig. 10A-10B). Finally, we determined the relative synaptic density from a given cone via its postsynaptic bipolar cells to each identified GC type (see details in *SI Appendix*, Figs. S9 and S10; *SI Appendix*, Table 2).

**Fig. 10.**
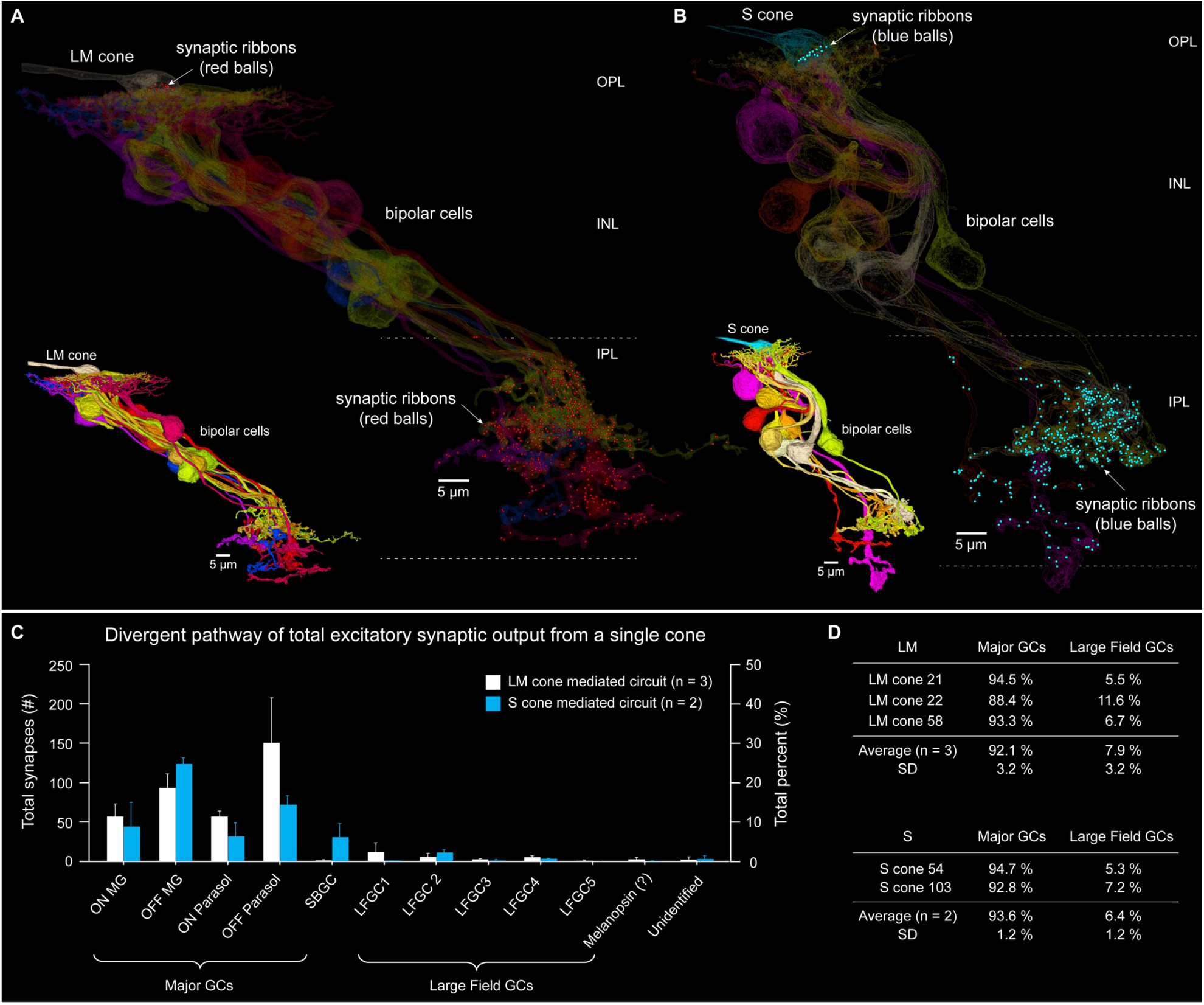
Vertical excitatory connectome. **A.** Vertical connectome for a single LM cone (cone 22; see details in *SI Appendix*, Fig. S9B). Cone pedicle and postsynaptic bipolar cells shown in silhouette. Synaptic ribbons in the cone pedicle (top, OPL) and bipolar cell axon terminals (lower, IPL) are shown as red balls. Inset shows all bipolar cells postsynaptic to this cone pedicle (outer, yellow-gold; inner, blue-violet-red). **B**. The same views for an identified S cone (cone 103; see details in *SI Appendix*, Fig. S10B). Synaptic ribbons shown as blue balls. **C**. Histogram shows output synapse numbers (left axis) and percentage (right axis) made by bipolar cells connected to 3 LM and 2 S cones to postsynaptic GC types. Note that OFF-pathway-derived synapses greatly outnumber ON pathway synapses. **D**. The great majority of synapses from LM and S cones are distributed to midget, parasol, and small bistratified ganglion cells (Major GCs), with all other large-field types receiving 7.9% (LM) and 6.4% (S) of the synaptic output (see also *SI Appendix*, Table 2). Unidentified processes account for 0.5% of the total synaptic output (LM) (see also *SI Appendix*, Table 2A-2C).

We found that bipolar cells that were postsynaptic to the three LM cones provided their output overwhelmingly to midget and parasol ganglion cells (*SI Appendix*, Fig. S9), accounting for 92.1% of the total bipolar synaptic output to ganglion cells (Fig.10C, 10D; *SI Appendix*, Fig. S9; *SI Appendix*, Table 2A-2C). The remaining ∼7.9% of bipolar synapses were directed to the large-field ganglion cell types (LFGC1-LFGC5, Fig.10C, 10D; *SI Appendix*, Fig. S9; *SI Appendix*, Table 2A-2C). Output to dendritic fragments of unidentified ganglion cells was rare, accounting for only ∼0.5% of the total bipolar output to ganglion cells (0.53 ± 0.76%, n = 3 LM cones; *SI Appendix*, Table 2A-2C). These data support the conclusion that, with the possible exception of extremely large-field and low-density melanopsin-expressing ganglion cells, we have accounted for all ganglion cell types present in our foveal sample.

## Discussion

### A human foveal connectome

Our aim was to provide a draft nanometer-scale connectome in the human CNS. Compared to ongoing studies of a larger volume of human neocortex(18), our foveal volume has the advantage that foveal circuitry is miniaturized and functionally complete – from receptors to ganglion cell outputs. Moreover, all cell types in the HFseg1 volume are identified and can be related to anatomical, physiological, and transcriptomic studies of human and non-human primate retina (*SI Appendix*, Table 1).

### Cell type profiling

Comparable datasets to HFseg1 exist for primate central retina where cell types were clustered by gene expression profiles(12, 13, 15, 43, 88). The HFseg1 volume permits a comparison of the two approaches and suggests generally good correspondence, though with some unexpected differences. For example, the ratio of H1 to H2 horizontal cell types identified by transcriptomic analysis was shown in the macaque and human central retina to be ∼3:1 and 7:1, respectively. By contrast, the H1 to H2 ratio in HFseg1 is much higher (22:1), with H1 cells comprising 96% of the horizontal cells at this foveal location. Thus, the trend toward H1 cell dominance in the transcriptomic dataset is evident but more extreme in HFseg1. A similar trend holds for other clearly identified types: for example, midget ganglion cells make up ∼90% of the ganglion cell population in our sample and 86% in the transcriptomic dataset; Aλλ amacrine cells make up 26% of the amacrine cells but only 18% of the amacrines identified by transcriptomic analysis. One explanation for these differences may be that our smaller sample represents the cone-enriched foveal center, whereas the larger samples used for transcriptomics encompassed a much larger area (∼1.5 mm, central 5 degrees). Cell type-specific density changes are extremely steep across this region, thus, larger spatial samples would average out such changes.

Certain cell populations previously recognized were not evident in HFseg1. For example, the recent functional, anatomical, and molecular distinction made between DB3a and DB3b in the macaque monkey(79, 89) and human retina(13, 38) was not apparent in HFseg1. Further, we found evidence for only 19 amacrine cell types, whereas molecular profiling in human retina (including the retinal periphery) has proposed 73 amacrine cell types(15). By comparison, 63 amacrine clusters have been identified in mouse retina(90, 91). These differences suggest that the relationship between cells defined by gene expression and anatomical types may not always be straightforward. A further implication is that the extreme increase in density of the midget circuit, required for high acuity and color vision, gives rise to an attendant density increase in other foveal cell types tightly linked to the midget pathway (e.g., H1 horizontal cells).

### The vertical excitatory connectome

In the non-human primates, fewer than 20 ganglion cell types have been defined, with transcriptomic and anatomical identification in close agreement(12, 32, 33). In the human fovea, only 12 pathways have been suggested by transcriptomic analysis(13, 15). It was nevertheless striking that in the HFseg1 dataset (*SI Appendix*, Table 1H), ∼96% of the ganglion cells comprised just five anatomically distinct pathways: the ON and OFF midget and parasol cells, and the small bistratified cells. When we manually traced the vertical connectome from a single cone to its postsynaptic bipolar and ganglion cells, we found that ∼92% of synaptic output targeted these same five pathways (Fig. 10D), confirming their dominance. Thus, a fundamental, defining feature of foveal synaptic organization may be a reduction in the number of visual pathways compared to the peripheral retina.

Over 40 distinct ganglion cell populations have been recognized in the mouse retina, with molecular profiling, physiology, and anatomy in agreement(29–31). What explains this dramatic reduction in ganglion cell types from the mouse retina to the primate fovea? The difference suggests that humans and mice have evolved fundamentally different solutions to analyzing the visual scene. Great emphasis has been placed on the behaviorally relevant feature-detection properties of diverse mouse ganglion cell types(92–94). By contrast, in primates, considerable emphasis has been placed on the dual and more general role of the midget pathway in both form and color perception(74, 95, 96).

Recent molecular orthotype analysis shows that the midget and parasol ganglion cell types are likely homologous to sustained and transient alpha cell types, respectively, in the mouse(14). What differs dramatically, however, is the extremely large dendritic tree and consequent low spatial density (∼2% of the total ganglion cells) of the mouse ortholog of the midget ganglion cell. Thus, the mouse retina sacrifices spatial resolution(97), opting instead for an expansion of feature detectors that mediate visually driven reflexive behaviors—with mice employing feature-detection strategies analogous to those described in classical studies of amphibian vision(98). In primates, the midget circuit represents an evolutionary extreme where the spatial resolution afforded by tightly packed photoreceptors is preserved through private-line connectivity. This high-resolution sampling is behaviorally exploited through active vision strategies. Humans not only explore a visual scene by making saccadic eye movements but also make small fixational eye movements (microsaccades) to extract information at peak resolution(99). These microsaccades are dynamically modulated during highly precise manipulation (e.g., threading a needle)(99–101). Thus, the incessantly mobile fovea and related retinal circuits are designed for inspection, manipulation, interpretation, and understanding of an object—the converse of a feature detector.

### The foveal connectome and human color vision

*S cone input to H1 cells.* Most psychophysically based models of human color vision assume the presence of two parallel color-opponent channels: red vs green and blue vs yellow^109^. The red-green pathway shows opposing L vs M cone interaction and has been associated with the circuitry that feeds midget ganglion cells(102). However, the human psychophysically defined red-green pathway includes a small S cone input(103). The locus of this critical signal pathway remains unclear, with recent data from the macaque monkey implicating both the retina^110^ and primary visual cortex^111^. The HFseg1 connectome shows that H1 horizontal cells, a major source of the surround in the midget circuit^51^, likely contribute antagonistic signals from all three cone types (L, M, and S) to the midget pathway. In this scenario, the human foveal midget circuit would derive its center response from either an L or M cone, while the surround would provide cone opponency by non-selectively contacting all three cone types(73, 104, 105), with the S cones making a small contribution to the chromatic receptive field, creating L+S vs M or M+S vs L opponency.

*S-LM cone coupling.* As a first step in deriving the foveal connectome, we determined the “contactome” among the tightly packed cone synaptic pedicles. Contacts among the cone types harbor gap junctions and thus form electrical synaptic connections between cones(65, 69). We confirmed and extended previous observations suggesting that, in contrast to non-human primates and non-primate mammals, human foveal S cones show extensive telodendritic contacts with neighboring LM cones(44, 68), and these contacts are identical in form to LM-LM cone contacts.

A network model of this connectivity showed that, as a previous model suggested(66), mixing S and LM cone signals can alter the action spectrum of the S cone light response, with increased sensitivity at longer wavelengths. This result raises a functional puzzle: the prevailing view is that electrical coupling serves to increase signal relative to noise by reducing the uncorrelated voltage difference (i.e., the noise component) among neighboring cones and thereby increase contrast sensitivity(66). However, any mixing of S and LM cone signals would detract from the separation of cone signals required for color processing, in turn raising the question of why such coupling occurs in human foveal retina, whereas it is absent in both the macaque and marmoset retina(44).

The answer may be related, at least in part, to the connectivity of H1 horizontal cells with S cones noted above. Coupling among the S, L, and M cones prior to this connection may serve to preserve the smaller S cone signal within the H1 cell network. This in turn may be critical for the unique requirements of human color vision whereby a small S cone signal contributes critically to the “red-green” chromatic channel defined psychophysically(103). This hypothesis could be tested by adding the horizontal cell connectome to the current cone-cone network and testing the effects of noise and coupling on the spectral tuning of the H1 cell response and ultimately its contribution to cone opponency modeled in the midget ganglion cell.

## Acknowledgements

This work was supported by NIH grants EY-028282 and RF1-MH129260 to D.M.D. and NIH grant P51 OD010425 to the Washington National Primate Research Center and EY01730 to the Vision Research Core at the University of Washington. We thank Sharm Knecht for tissue preparation and managing EM data acquisition.

## Author contributions

Y.J.K. and D.M.D. conceived the project. Y.J.K. and D.M.D. generated the ground truth for segmentation and synapses, performed proofreading and created annotations. R.G.S constructed and ran the computational models. Y.J.K. and D.M.D. analyzed the data and generated figures. Y.J.K., D.M.D., P.R.M., and U.G., wrote the paper and consulted on cell type annotations and figure development. C.A.C. and R.G.S., edited the paper. A.P. acquired human tissue. D.M.D., Y.J.K., C.A.C. and A.P. acquired funding. Zetta AI performed EM data alignment, segmentation, synapse detection and cone-cone contactome detection and ingested all data into the CAVE infrastructure for proofreading. S.G. developed visualization and analysis tools at NeuroMaps.app.

## Declaration of interests

K.L., N.K., D.I., T.N., R.L., S.P., A.H., J.A.B., J.S. and T.M. declare financial interests in Zetta AI. S.G. is owner of Aware LLC and developed the NeuroMaps.app. The remaining authors declare no competing interests.

## Methods

### EXPERIMENTAL MODEL AND SUBJECT DETAILS

#### Human tissue acquisition and preparation

Human eyes (52-year-old, white male; brain dead organ donor) were acquired from the Medical University of Vienna, Vienna, Austria at the time of death by surgical enucleation. A small incision was made to give fixative access to the posterior chamber and the retina was in this way immersion fixed in 4% glutaraldehyde in 0.1 M sodium cacodylate buffer, pH7.3-7.4 at room temperature. Medical history confirmed no abnormalities of the visual system recorded and no driving limitations; the ocular status was defined medically as unremarkable at time of enucleation. Eyes were shipped in fixative to the University of Washington where the retinas were dissected and prepared for electron microscopy.

### METHOD DETAILS

#### Identification of foveal ROI, serial block-face SEM sample preparation and image acquisition

Retinal tissue was dissected under a dissecting microscope to facilitate identification of the optic disc and the center of the foveal pit as well as the major retinal meridians. We selected an ROI that extends across absolute eccentricity of ∼450-650 microns (temporal retina). This location is linked to cones located ∼100-300 microns from the foveal pit center due to the lateral migration of retinal circuits from the cones at the foveal center. This location encompasses much of the central 1 degree (∼300 µm/deg) of visual angle (Fig. 1A-1C).

The dissected retina was then plastic embedded and prepared for electron microscopy as previously described^1^. In brief, the tissue was incubated in a 1.5% potassium ferrocyanide and 2% osmium tetroxide (OsO4) solution for 1 h. After washing, the tissue was placed in a freshly made thiocarbohydrazide solution for 20 min and then rinsed and incubated in 2% OsO4 for 30 min. Lastly, the tissue was stained en bloc in 1% uranyl acetate overnight and subsequently stained with Walton’s lead aspartate, dehydrated and plastic embedded (Durcupan, 44610, Sigma Aldrich). After evaluation of the ROI by semithin sectioning the tissue block was trimmed, gold-coated by standard methods, and mounted in an Apreo SEM Volumescope (Thermo Fisher). We cut 3028 sections at 50 nm thick for a total depth of ∼150 microns, imaging an ROI (180 µm tall x 180 µm wide; 5 nm pixels) that encompassed the full vertical depth of the retina, from the Henle fiber layer (HFL) to the nerve fiber layer (Fig. 1D). The overall ROI was imaged on the blockface after each section as 25 sequential tiles (8000 x 8000 pixels or 40 x 40 µm; 75,700 total image tiles) with 10% overlap. Images were stitched within layer and aligned across layers using standard methods available in TrakEM2^2^ (NIH image plugin; Scale Invariant Feature Transform (SIFT) with affine translation) to create a good quality volume. We first evaluated this TrakEM2-aligned volume by manual reconstruction of a small number of selected cells and circuits(106). To further develop this volume as a connectomic resource we applied deep-learning based computational approaches to realign, segment and reconstruct all cells and synaptic connections as follows(20, 39).

#### Volume creation

*Montaging.* Tile offsets were determined manually, then refined by template matching at 40 nm with a search window of 96 pixels. Tiles were cropped by 128 pixels to avoid edge distortions and placed to minimize the sum of least squares from the new offsets. Outliers were inspected and given manual placement. These cropped tiles were processed to create a pyramid of encodings for resolutions between 20 nm and 640 nm(107), and then stitched to make the montaged section in two steps. In the first step, only the two neighboring tiles in Y were considered, with both neighbors being fixed. Using an online finetuner pyramid(107) with the calculated placement as the starting point, a displacement field was generated that would warp the encoded tiles to minimize the pixel difference with the fixed tiles.

The encoded tiles were then warped using this displacement field, eroded by a pixel to avoid partial pixels at the edge, blended to the fixed tiles using the mean of the pixel values in the overlap, and then passed through CLAHE (Contrast Limited Adaptive Histogram Equalization) enhancement to mitigate the contrast loss from the warping and blending, resulting in strips of encoded tiles where every other tile had been warped. In the second step, the strips of tiles from the first step were stitched to each other, holding every other strip fixed in a similar process. An error map was generated at both steps by computing the difference between the encodings in the overlap at 40 nm and inspected. The non-encoded tiles were cropped by 394 pixels, warped according to the composed displacement fields from the encodings, and combined to yield montaged sections.

*Alignment.* The unaligned sections were first processed to create a pyramid of encodings(107). A rigid transformation for the entire dataset was estimated using SIFT features(108) extracted from the encodings at 640 nm and correspondences established between nearest and next-nearest section pairs. From the rigidly aligned encodings, displacement fields at 40 nm resolution were generated between nearest and next-nearest section pairs using an online finetuner pyramid(107).

A convolutional net trained to detect local misalignments and/or simple mismatches caused by defects in the EM images was used to mask unreliable regions in each displacement field. In addition, 25 sections with defects that presented with non-smooth deformations were masked by hand. Pairwise displacement fields and transformation masks were then passed to a global relaxation(107, 109). This relaxation was performed first at 2560 nm across all 3028 sections of the dataset, and subsequently in blocks of approximately 100 sections at a resolution of 640 nm, using the previously generated drift-free fields as constraints for the start and end sections of each of the 31 blocks. The resulting displacement fields were composed with the earlier rigid transformation and the final aligned image stack warped and rendered at 10 nm.

*Segmentation.* We pretrained a convolutional net to predict affinities between neighboring voxels at 16×16×40 nm³ using 1,368 µm³ of manually annotated data derived from three mouse visual cortex datasets(110). We used the residual symmetric U-Net architecture(111) with modifications(112) and trained with data augmentation previously described (113). The network was trained for 2,000,000 iterations using four NVIDIA L4 GPUs with <u>PyTorch</u> data parallelism, where each GPU processed individual patches with its own model replica.

To further fine-tune this network, we manually created training data consisting of 39 µm³ at a resolution of 20×20×50 nm³. In contrast to the pretraining datasets, mitochondria were intentionally over segmented to bias boundary detection toward a more conservative approach, aiming primarily to reduce merge errors at the expense of increasing split errors. We fine-tuned the network for 300,000 iterations.

We predicted an affinity map for the whole dataset. Dark voxels could sometimes span multiple cells and cause segmentation errors. We therefore applied a binarized mask for dark voxels with a threshold of 0.2, invalidating each dark voxel and its three incident affinities. The resulting masked affinity map was then segmented with a mean affinity agglomeration threshold of 0.25(113). The segmentation was ingested into the Connectome Annotation Versioning Engine proofreading platform, CAVE(114) for further analysis.

CAVE permitted proofreading to be performed by a user community accessing the web-based neuroglancer interface(113, 115) with dynamic updating of segment ID and related synaptic connectivity. The HFseg 1 segmentation was of high initial quality such that it was possible to identify and annotate all major cell classes within the volume and to sort these cells into populations of distinct types using the annotation tools available in neuroglancer. All cells are accessible at NeuroMaps (https://neuromaps.app).

Cell types were identified provisionally based on segment 3D morphology (dendritic morphology, stratification depth, and spatial regularity and interaction (mosaics) and in some cases nuclear morphology). Four of the authors (Kim, Dacey, Grünert and Martin) conferred and agreed on cell type identification. Proofreading is ongoing (∼ 40,000 edits) and applied thus far primarily to correct larger obvious mergers (primarily with glial cell processes) that interfered with cell type identification and additional more detailed proofreading on circuits of interest in the current survey of cone-bipolar-ganglion cell connections. Many cell populations were unambiguously recognized based on previous studies of primate retina(41, 116, 117) (e.g., the S cones, rods and LM cones, the H1 and H2 horizontal cells, the starburst and Aλλ amacrine cell types, the midget, diffuse, blue-cone bipolar, and rod bipolar types, or the midget, parasol and small bistratified and large bistratified ganglion cell types; Müller glia, astrocytes and microglia); some cell types (the low-density amacrine types) were provisionally designated subject to further proofreading and connectivity analysis. Following FlyWire(40), we have initiated an open community of retinal neurobiologists (currently 7 labs participating) to use these tools to complete proofreading of the HFseg1 volume.

#### Synapse detection

*Ribbon synapse detection in OPL and IPL.* A symmetric residual U-Net architecture was trained on data with a resolution of 10×10×50 nm^3^ using input patches of size 128×128×20 to generate a probability map that indicates the likelihood of each voxel belonging to a synaptic ribbon (118). To remove less robust predictions at the boundaries, only the central 96×96×16 voxels of the input patch was used as the output. The model was trained on a total volume of 2,254 µm^3^ of manually labeled cutouts sampled from various regions across the entire dataset.

The ribbon segmentation was produced by (1) down sampling the probability map by 2×, (2) thresholding for voxels above 0.15, and (3) using 26-connectivity to extract connected components. Segments smaller than 5 voxels in a 20×20×50 nm^3^ resolution were neglected to eliminate spurious detections.

To achieve ribbon detection in the OPL segments smaller than 5 voxels (in a 20×20×50 nm^3^ resolution volume) were first disregarded to eliminate spurious detections of extremely small ribbon-like structures. For each detected ribbon within a cone pedicle, postsynaptic partners were identified via the following steps: (1) All neighboring segments touching the pedicle within 800 nm of the ribbon centroid were identified. (2) For each neighbor, a representative point close to the ribbon was chosen as the postsynaptic site. These constitute the set of putative synapses for each ribbon. (3) After cell identification, only putative synapses onto bipolar or horizontal cells were kept; all others were filtered out (e.g. Müller glia, neighboring cones, mitochondria). (4) Finally, for each distinct ribbon-postsynaptic cell pair, all but the shortest putative synapse were filtered out. The result was that a single synaptic connection was found onto any bipolar or horizontal cell contacting that cone pedicle within 800 nm of a given cone ribbon.

For synaptic ribbons in the IPL, segments smaller than 5 voxels in a 20×20×50 nm^3^ resolution within the bounds of the IPL were also disregarded to eliminate spurious detections. For each detected ribbon, postsynaptic partners were automatically identified via the following steps: (1) Identify the two largest cells within a small (500 nm) window of the centroid of the ribbon which are adjacent to the presynaptic cell and are also adjacent to each other. (2) If such a pair is found, the ribbon is assumed to be presynaptic to both (i.e., a dyadic synapse). (3) If no such pair is found, then within the window, the single largest profile adjacent to the presynaptic cell is identified as the postsynaptic partner. (4) Putative synaptic connections were filtered by cell type, requiring the presynaptic cell to be previously annotated as a bipolar cell, and the postsynaptic partner to be previously annotated as amacrine, or ganglion cell. Note we did not at this time try to use a convolutional net to detect a low density of possibly ribbonless synapses.

*Vesicle cloud detection and conventional synapse assignment in the IPL.* A symmetric residual U-Net architecture was trained on data with a resolution of 10×10×50 nm^3^ using input patches of size 128×128×20 to generate a probability map that indicates the likelihood of each voxel belonging to a vesicle cloud(118). To remove less robust predictions at the boundaries, only the central 96×96×16 voxels of the input patch was used as the output. The model was trained on a total volume of 2,254 µm^3^ of manually labeled cutouts sampled from various regions across the entire dataset (the same cutouts as ribbon synapse detection). Manual labels were roughly annotated to mark approximate locations of vesicle clouds rather than precise boundaries.

The vesicle cloud segmentation was produced by (1) downsampling the probability map by 2×, (2) thresholding for voxels above 0.15, and (3) using 26-connectivity to extract connected components. Segments with fewer than 200 voxels were excluded, based on the range of vesicle cloud sizes encountered.

*IPL conventional synapse assignment*. A symmetric residual U-Net architecture was trained on data with a resolution of 20×20×50 nm^3^ using input patches of size 24×24×8 to generate a probability map that indicates the likelihood of each voxel belonging to the postsynaptic cell(118). The model was trained on 682 µm^3^ of manually annotated labeled cutouts.

The model was applied once for each vesicle cloud detected, in a window centered on the centroid of the vesicle cloud. Using the initial automatic cell segmentation, the cell with the highest mean probability within the window was selected as the putative postsynaptic partner(118). Further filtering by cell type was applied, requiring the presynaptic cell to be previously annotated as an amacrine cell, and the postsynaptic partner to be previously annotated as amacrine, bipolar, or ganglion cell.

#### Building and modeling the cone-cone contactome

*Generating the cone-cone contactome.* We identified gap junctions between cone pedicles and between cone pedicles and rods as follows. Cones and rods were manually identified and proofread to completion. Contacts were computed using a recently developed supervoxel agglomeration process (abiss)(119) where a contact is considered to be a set of connected voxel faces that are shared by the same pair of segments. For each contact, we counted the number of voxel faces along each orientation (xy, xz, and yz), as well as the minimum bounding box containing all of the faces. The surface area of a contact was estimated by computing the sum of the area of all faces involved. Contacts were excluded from analysis if (1) the surface area of the contact was ≤0.04 μm^2^ (mean surface area of contact, 0.55 ± 0.3 µm^2^, n = 1141 cone-cone contacts) or (2) the center of the bounding box was more than 15 μm from the approximate center of the cone pedicle (manually identified). All remaining contacts were manually inspected with 1× coverage to determine if they corresponded to what would be manually identified as a true cone-cone contacts harboring gap junctions.

*Biophysical modeling of cone-cone contactome*. To generate a biophysical model of the electrically coupled photoreceptor network, we started with data from the X,Y locations of 310 cone and 24 rod synaptic terminals, and for the cones their M/L vs. S identity. We then incorporated all reviewed and accepted cone-cone contacts and the surface areas of each contact.

The cone morphology was modeled as a sphere (single compartment, dia. 2.5 µm) for the outer and inner segments and soma, connected to an axon (dia. 1.6 µm, Ri 80 Ohm-cm, Rm 12,000 Ohm-cm^2^ (120); V_rev_ −60 mV, compartment size 0.05 lambda, ∼9 compartments/cone axon) extending 300 µm to the terminal (diam 5 µm, Rm 12,000 Ohm-cm^2^, V_rev_ −60mV). Rod morphology was similar (OS/IS diam 2.5 µm, axon dia 0.45 µm, terminal dia. 3 µm). The OS/IS/soma compartment contained a transduction mechanism (max conductance 880 pS) with spectral sensitivity appropriate for the cone identity. The transduction mechanism of the photoreceptor biochemical cascade generated a peak ∼20 ms after the stimulus with a damped recovery (121–128). Spectral sensitivity for the L/M cones was taken from macaque monkey (128) and for the S cones from human cone data (peak sens. 420 nm (129) corrected for lens spectral sensitivity (130)). In some models including both lens and macular filters, the S cone peak sensitivity was shifted to ∼440 nm.

The location of all the S-cones and rods in the reconstructed contactome was known but the individual identity of L/M cones was unknown, so for the simulations the L/M identity was set randomly. The target L/M ratio was 1.7 and the actual L/M ratio in the model shown was 1.697. The contacts between cones had not been resolved into gap junctions (groups of connnexin molecules), but the area of each contact had been measured. The conductance for each contact was calculated by taking the average area of the contacts (0.55 µm^2^), assuming 5 connexins (69) (15 pS per connexin) per average contact (131, 132) to produce an average conductance of 75 pS, or 136 pS/µm^2^. The conductance for each individual contact was then calculated by multiplying by the contact area. Contacts were simulated with a linear gap junction conductance (i.e. electrical synapse). The cones had on average 3.6 contacts with neighboring cones. The average L/M total coupling conductance was ∼270 pS, and for S-LM cones was ∼282 pS. There were a total of 715 coupled pairs of cones, 659 LM pairs, and 56 pairs with 1 S cone. The average coupling conductance between adjacent pairs of L/M cones was 116 pS, and for L/M–S pairs was 123 pS. However, the average measured coupling conductance between LM pairs was 133 pS (n = 25) and between S-LM pairs was 166 pS (n = 14) due to the existence of multiple current paths between pairs through the gap junction network. The average cone terminal input resistance was ∼0.4 Gohm.

The simulated cones were stimulated with flashes of monochromatic light (10 ms, 1000 µm spot, ∼1 x 10^5^ photons/µm^2^/sec) on a white background (1 x 10^4^ photons/µm^2^/s, spectrally flat). The resting potential V_rest_ in response to the background was ∼ −40 mV but varied by ∼0.3mV depending on the cone identity and coupling. The flash response varied from 0 to −5 mV. To determine the effect of the gap junctions, simulations were run with the gap junction conductance at 0%, 50%, 100%, and multiples of the nominal 136pS/µm^2^ conductance. The model was constructed in the Neuron-C simulation language (133) and comprised ∼3000 compartments. The model was run on AMD Opteron 3000 MHz CPUs under the Linux and Mosix operating systems. Approximately 200 simulations were run of the 310-cone model to develop and calibrate the model. Modeling code can be accessed at zenodo.org upload of Neuron-C at: https://doi.org/10.5281/zenodo.17792059.

#### Proofreading, synaptic connectivity and updating with the Connectome Annotation Versioning Engine (CAVE) and NeuroMaps

As noted above we ingested synapse predictions and cell type annotations into CAVE. A critical functionality given by CAVE is that is allows collaborative users to programmatically query the up-to-date proofread connectivity graph (synaptic connectivity is updated during proofreading of each segment-cell type) as well as any changes made to the metadata (cell type IDs, descriptions, or other annotations). We then adapted NeuroMaps to combine this metadata with synaptic connectivity for each segment ID and to permit synaptic proofreading (addition or deletion of specific synaptic predictions).

## DATA AND SOFTWARE AVAILABILITY

The HFseg1 dataset (volume, segmentation, synaptic detection and full annotation) are currently hosted by Zetta.ai (https://zetta.ai/) and all data is accessible for community proofreading by permission at NeuroMaps. The latest version of the HFseg1 dataset is also available for interrogation at NeuroMaps (https://neuromaps.app). Modeling code can be accessed at zenodo.org upload of Neuron-C at: https://doi.org/10.5281/zenodo.17792059

## Author Contributions

Y.J.K. and D.M.D. conceived the project. Y.J.K. and D.M.D. generated the ground truth for segmentation and synapses, performed proofreading and created annotations. R.G.S constructed and ran the computational models. Y.J.K. and D.M.D. analyzed the data and generated figures. Y.J.K., D.M.D., P.R.M., and U.G., wrote the paper and consulted on cell type annotations and figure development. C.A.C. and R.G.S., edited the paper. A.P. acquired human tissue. D.M.D., Y.J.K., C.A.C. and A.P. acquired funding. Zetta AI performed EM data alignment, segmentation, synapse detection and cone-cone contactome detection and ingested all data into the CAVE infrastructure for proofreading. S.G. developed 3D visualizations and analysis tools at NeuroMaps.app.

## Competing Interest Statement

The authors declare no conflict of interests.

## Notes

https://neuromaps.app/datasets

### Summary of Updates

We have revised our manuscript for submission to a journal.

