## Supporting Information for "Connectome of a human foveal retina"

**This PDF file includes:**

Supporting text
Figures S1 to S10
Tables S1 to S4
SI References

### Supporting Information Text

#### Cell type identification

**Rods and cones.** Rod and cone photoreceptors were distinguished by the well-established and distinctive morphology of their axon terminals, the small rod spherules (n = 24) and the large cone pedicles (n = 313). We also confirmed that the invaginating contacts to the rod spherules derived from subsequently identified rod bipolar cells. Note that rods could also be distinguished from cones by the much smaller diameter of their axons within the Henle fiber layer (HFL). These features were not quantified but are evident in the segmented 3D view of the cell's morphology.

Unlike in macaque and marmoset retina, human S cone pedicles (n = 17) were not easily distinguished by smaller overall size, reduced number of synaptic ribbons or lack of telodendritic contacts with neighboring cone pedicles (Fig. 2A; *SI Appendix*, Table 1C). We therefore relied on the distinctive dendritic and axonal morphology of the blue cone bipolar type (BB) that makes selective invaginating contact with S cones and the concomitant lack of an ON-midget bipolar (IMB) connection. The L and M cone pedicles have not been previously distinguished by morphology or connectivity, with one possible exception[1]. In the present dataset we have not yet attempted to determine if two groups of non-S cone pedicles are present and have used LM to refer to the combined L and M cone pedicles (n = 296) (Fig. 2A; *SI Appendix*, Table 1C).

**Horizontal cells.** Morphological and connectional differences defining H1 and H2 horizontal cell types in the primate retina are well established and largely consistent in the HFseg1 volume[2, 3, 4, 5]. However, H1 cells frequently contact S cones and greatly outnumbered H2 cells. The H1 cells (n = 256; *SI Appendix*, Table 1E) were distinguished by small (~30–50 μm diameter) and profusely branched dendritic trees (Fig. 3A). Neighboring H1 cells overlapped extensively such that each cone pedicle was contacted by several H1 cells. A characteristic single, long axon-like process arose from the soma and extended to the edge of the volume within the OPL without branching. These long unbranched processes are known to terminate in a profuse arbor that innervates rod spherules[6, 7]. All H1 cell axons in HFseg1 extended beyond the volume boundaries and did not contact the rod spherules within the volume, which were therefore innervated by axonal processes that originated from H1 cells outside the volume. In contrast to H1 cells, the H2 cells (n = 12; *SI Appendix*, Table 1E) showed very large, and loosely branched dendritic trees (>100 μm in diameter) that showed little dendritic overlap with neighboring H2 cells[2, 8, 9] (Fig. 3B). The long H2 cell dendrites tended to converge on and contact S cones located within their fields, while only sparsely contacting LM cones (*SI Appendix*, Fig. S4). The axon of the H2 cell is distinguished from that of the H1 cell in that it takes a meandering course, branches sparsely, and gives rise to lateral elements in widely spaced cones, most of which are S cones.

**Bipolar cells.** In foveate primates, flat (FMB, OFF type) and invaginating (IMB, ON type) midget bipolar cells show dominant connections to single cones and ganglion cells, whereas diffuse types contact several cones and comprise eight previously recognized types (DB1, 2, 3a, 3b, 4, 5, 6 and Giant)[10] (Fig. 4). A single blue cone bipolar (BB) cell type shows selective targeting of S cones (*SI Appendix*, Fig. S5), and a single rod bipolar type makes selective contact with rod spherules. It was possible to recognize and annotate all of these types in HFseg1 with the exception of the DB3a-DB3b distinction recently made in macaque and human retina[11, 12, 13, 14, 15, 16] and the possible absence of the Giant type.

Of the cells with distinctive cone connectivity, FMB (n = 302; *SI Appendix*, Table 1F) and IMB cells (264 cells; *SI Appendix*, Table 1F; Fig. 4A, 4B) accounted for 62% of all bipolar cells. Blue-cone bipolar cells (BB cells; Fig. 4B, 4C; *SI Appendix*, Fig. S5) by contrast formed only 1.9% of all bipolar cells (17 cells; *SI Appendix*, Table 1F). Of the presumed OFF diffuse bipolar types, we identified DB1[17] (54 cells), DB2 (92 cells) and DB3 (41 cells; *SI Appendix*, Table 1F; Fig. 4A, 4C). We could not further divide the outer DB3 cells into previously identified DB3a, DB3b types[11, 12, 13, 14]. Instead, we found only two bipolar populations that costratified in the IPL and tiled the IPL with their axonal arbors that we therefore referred to as DB2 and DB3. The DB3 cells showed large axonal fields and likely correspond to the calbindin-positive DB3a type in macaque, marmoset and human retina[15, 16, 18, 19]. If this association holds then DB3b cells are either absent or present at a very low density in the foveal DB population. It remains possible that a detailed analysis of synaptic connectivity for the OFF-DB cells might reveal a distinct DB3b population. However, the clear mosaic organization of each type as currently described would make this possibility less likely.

In the inner half of the IPL, DB4 (n = 34), DB5 (n = 45) and DB6 cells (n = 12, *SI Appendix*, Table 1F; Fig. 4) were distinguished by axonal morphology, non-overlapping spatial arrangement and/or stratification depth in the IPL. A small number of cells that remain to be studied in detail could include the Giant type (n = 7, *SI Appendix*, Table 1F). Lastly, rod bipolar cells (RB, n= 6 cells, 0.7% of total BCs, *SI Appendix*, Table 1F) made invaginating contacts[20, 21], with rod spherules and showed small unbranched axonal arbors that contained far fewer synaptic ribbons (15.8 ± 2.2 ribbons, n = 5) compared to diffuse or midget bipolar cells (see the detailed number of synapses in *SI Appendix*, Fig. S9)[22, 23]. As presented in the Results, x-type bipolar cells (n = 20; *SI Appendix*, Table 1F) were further distinguished by a lack of an outward extending dendrite but the presence of abundant synaptic ribbons in the axonal arbor (see also *SI Appendix*, Fig. S6). DBbroad cells (n = 18; *SI Appendix*, Table 1F) were distinguished by an axonal arbor that extended variably into both the outer-OFF and inner-ON IPL and were presynaptic to both ON and OFF ganglion cell types. In the OPL, the DBbroad cell dendrites remain to be completely proofread, but thus far we found only basal contacts with cone pedicles and have placed the DBbroad cells provisionally in the OFF bipolar category, though the lack of invaginating cone contacts does not preclude an ON-type response from these cells[24].

**Amacrine cells.** Amacrine cell types show great morphological diversity and have been divided into many more types than other retinal cell classes. The result is that, with a few well-studied exceptions, there is little consensus on the number of amacrine cell types and how they compare across species. To annotate HFseg1 amacrines we started with the high-density, small field AII amacrine (AC2, n = 75 cells; *SI Appendix*, Table 1G) that surprisingly accounted for over 25% of the amacrines with cell bodies in the volume. These cells showed the characteristic lobular appendages making synaptic output to cone bipolar cell axon terminals and long arboreal dendrites stratified near the IPL-GCL border, despite the lack of rod bipolar axon terminals at this foveal location (Figs. 5A, 7C and 9). We divided all small field (< 100 μm diam) amacrine cells, including ACII, into six provisional types (AC2, 3, 4, 5, 6, and 11) based on dendritic morphology, spatial tiling and IPL stratification depth, pending a more detailed analysis of their synaptic connectivity. Typical of the small-field types, AC11 illustrated in Figure 5 (n = 22; *SI Appendix*, Table 1G) showed fine, densely branched, dendrites that define a group of small field cells conventionally referred to as “knotty” amacrines[25, 26, 27]. The broadly stratified AC11 may correspond to the strongly parvalbumin-positive cells described in macaque monkey[28]. Small-field cells together accounted for ~65% of the amacrine cells in HFseg1.

Cells with larger (> 100 μm) dendritic fields were divided into 13 provisional types based mainly on morphology (including nuclear staining pattern; see *SI Appendix*, Fig. S3) and stratification depth in the IPL. Some of these types were easily classified by well documented morphology (e.g., the starburst amacrine cells illustrated in Fig. 5 or the interplexiform cells illustrated in *SI Appendix*, Fig. S8). Almost all starbursts were found in the ganglion cell layer (GCL; see Fig. 5D), where they were the most numerous AC type (14/40 cells, 35%), consistent with previous estimates[29, 30]. Other large-field types like AC12 (n = 6; *SI Appendix*, Fig. S8; *SI Appendix*, Table 1G) or AC8 (n = 10 cells; Fig. 5G, 5H; *SI Appendix*, Table 1G) showed a stereotyped morphology and stratification pattern that made them relatively easy to classify. Several large-field types showed sparsely branching dendrites and varied stratification patterns; for these groups, an analysis of synaptic connectivity will be important for confirming or extending the current amacrine cell classification.

**Ganglion cells.** The morphology of the major, relatively high-density, ganglion cell types—the midget, parasol and small bistratified cells—is well established, and we therefore were able to annotate these clearly recognized types unequivocally without the need for detailed proofreading. As reviewed in the Results, these cells accounted for ~96% of the total ganglion cells with cell bodies within the HFseg1 volume, leaving only 25 ganglion cells that remained to be characterized. Parasol cells were distinguished by their well-established stratification depth, larger soma and dendritic field size relative to the midget ganglion cells which were unequivocally identified by their private or near-private line connection to midget bipolar cells. Small bistratified cells were distinguished by their relatively small but broadly stratified dendritic trees and dense input from blue-cone (BB) bipolar cells.

We divided the remaining large-field cells into 6 provisional types (five groups). Large-field type GC1 (LFGC1, n = 6 cells; *SI Appendix*, Table 1H; Fig. 6D, 6H) was postsynaptic to blue-cone bipolar cells and had a dendritic tree that co-stratified with the small bistratified cells. These cells likely correspond to the large bistratified GC observed in the human fovea[31] and marmoset and macaque retinal periphery[32, 33]. Consistent with input from blue-cone bipolar cells these cells showed an ON-response to S cone modulation in macaque retina[32]. Large field GC2 (LFGC2, n = 6 cells; *SI Appendix*, Table 1H; Fig. 6I, 6L) showed broadly stratified and densely branched dendritic trees; a counterpart in monkey retina remains unclear[34] but similar morphology has been observed in human peripheral retina[35]. A third cell group (LFGC3, n = 4 cells; *SI Appendix*, Table 1H; Fig. 6J, 6M) stratifies across the center of IPL (Fig. 6J, 6M) and appears to correspond to previously described recursive bistratified cells, identified as ON-OFF direction-selective cells in the macaque monkey retina[36]. A fourth cell group (LFGC4, n = 5 cells; *SI Appendix*, Table 1H; Fig. 6K, 6N) shows large cell bodies and radiating dendritic branching near the center of the IPL costratified with the inner-ON and outer-OFF parasol cells (three inner and two outer stratified types) and could correspond to the ON and OFF smooth mono-stratified types identified in macaque and marmoset retina[33, 36, 37]. A fifth group of cells shows very sparsely branching dendrites stratified in the inner IPL (LFGC5, n = 4 cells; *SI Appendix*, Table 1H) that could correspond to inner melanopsin cells[38] (or an inner large sparse type recognized in macaque and marmoset[33, 34]). Lastly, a few very sparsely branching GC processes (not linked to cell bodies within the volume) stratified along the outer border of the IPL were also present, likely corresponding to outer melanopsin ganglion cells[38]. If included in our current total, this would give seven large-field types and a total of 12 ganglion cell types in HFseg1.

**Glial cells.** The common radial glia of the retina, the Müller cells (MC, n = 555; Fig. 1; *SI Appendix*, Table 1B), were unequivocally recognized as a clear population of cell bodies in the approximate middle of the INL with large, irregularly shaped and euchromatic nuclei and a relatively filamentous, dark cytoplasm. A sparsely distributed population of glial cells was also present in the GCL (*SI Appendix*, Figs. S1B and S3B) and provisionally identified as astrocytes[39] (n = 32; *SI Appendix*, Table 1B). Astrocyte cell bodies and cytoplasm were similar in appearance to Müller cells. However, astrocyte cell bodies apposed blood vessels in the GCL (*SI Appendix*, Fig. S3B) and gave rise to multiple processes that extended radially into the GCL and IPL (*SI Appendix*, Fig. S1B) and tended to fasciculate along blood vessels. Processes of these astrocytes were restricted to the GCL and IPL and thus did not show the thick extension to the outer retina characteristic of Müller cells (*SI Appendix*, Fig. S1B). Microglial cells (n = 15; *SI Appendix*, Table 1B) had a characteristic elongated nucleus. Their morphology was distinct from both Müller cells and astrocytes (*SI Appendix*, Fig. S3A-S3C) with a very translucent cytoplasm containing apparent cellular debris. The density of microglial cells was low, consistent with previous measurements in the macaque monkey foveal retina[40, 41]. In the current volume, we have not yet attempted an analysis of blood vessels and associated pericytes, though these cellular elements are also segmented and are available to annotate and further characterize.

**Are there many large-field ganglion and amacrine cell types missing from our dataset?**

Both ganglion and amacrine cells include populations with large dendritic fields, and as discussed in the text concerning ganglion cells, we were alert to the possibility that some extremely low-density cells may be missed in our volume. For the ganglion cells, we used the vertical excitatory connectome to test the hypothesis that there were significant orphaned ganglion cell processes in the IPL that were not connected to their cell bodies of origin outside the volume. We did not find evidence of such processes, and indeed this is consistent with previous results showing that the large-field ganglion cell types in central retina show dendritic field diameters in the range of 100-150 µm. In addition, our anatomical picture is consistent with the connectomic dataset suggesting ~12 ganglion cell types (ON and OFF midget, ON and OFF parasol, small bistratified, LFGC1-LFGC5, and ON and OFF melanopsin fragments) in foveal retina. However, it is important to stress that our current analysis is ongoing and that synaptic connectivity may provide evidence for additional types.

Regarding amacrine cell types, we emphasize again that our current classification provides a provisional starting point for a deeper analysis. It is possible that detailed synaptic connectivity for all amacrine cells will lead to an increase (or decrease) in the number of anatomical-connectomic populations. It is also possible that some populations are so low in density that their cell bodies are not present in our current volume. As analysis of synaptic connectivity by amacrine cells proceeds, it should become clear if additional distinctive synaptic motifs are present that arise from longer, axon-like processes or very large-field amacrine cells. A complete analysis of synaptic connectivity should reveal the full complement of distinct amacrine populations, regardless of whether their cell bodies are present in the current volume. In addition, larger petascale volumes that encompass a greater area of the macula can address this question and also provide insight into how various cell populations change in density as the fovea is approached.

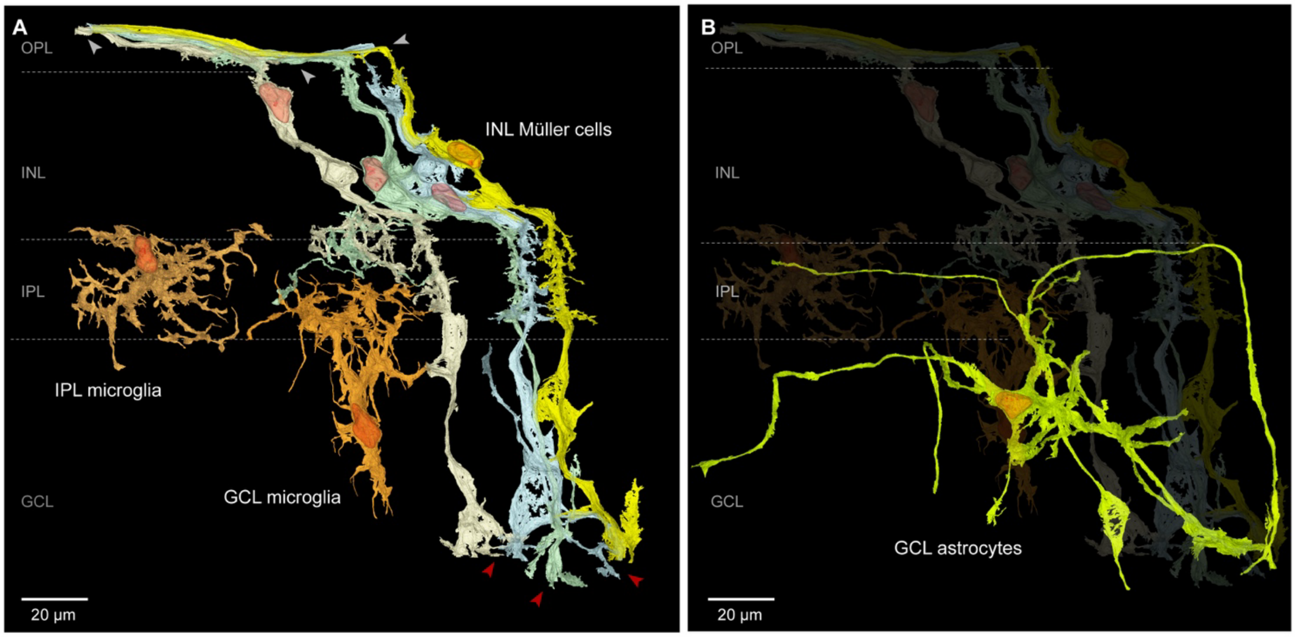


**Fig. S1. Non-neuronal cells. A.** Non-neuronal cells were divided into three types. Müller cells with cell bodies in the INL (INL Müller) were vertically oriented, extending outward processes into the Henle fiber layer ensheathing cone axons (white arrowheads) and inward processes to the inner surface of the retina (red arrowheads, Müller endfeet). A very small population of microglia (n = 15 cells) was present in the IPL with cell bodies near the INL-IPL border (IPL microglia; n = 10 cells) or in the GCL with processes that extended into the IPL (GCL microglia; n = 5 cells) and that showed a characteristic dendritic morphology and ultrastructure that distinguished them from Müller cells (see also *SI Appendix*, Fig. S3A-S3C). **B.** A third population of glial cells was present in the GCL (n = 26 cells; GCL astrocytes). Cell bodies of these cells were often apposed to blood vessels, and the long curving processes also tended to appose blood vessels, resulting in an overall stellate and tortuous branching pattern. The location of these cells in the GCL and their close relationship with the inner retinal vasculature suggest strongly that they correspond to astrocytes.

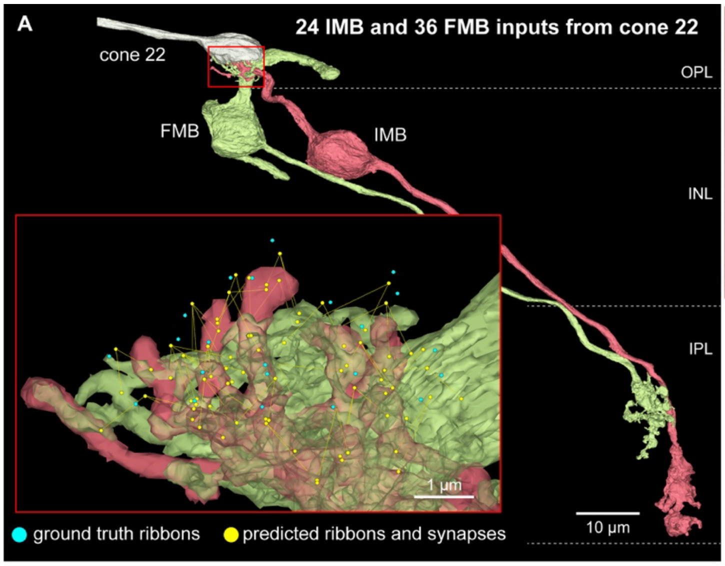


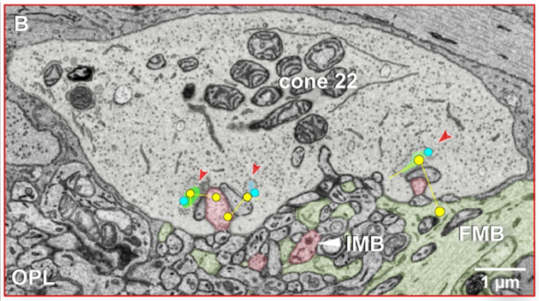


**C**

| LM | ribbon precision | ribbon sensitivity | ribbon total (#) | synapse precision | synapse sensitivity | synapse total (#) |
|---|---|---|---|---|---|---|
| cone 21 | 100 % | 100 % | 12 | 98 % | 100 % | 43 |
| cone 22 | 88 % | 100 % | 22 | 98 % | 99 % | 61 |
| cone 58 | 89 % | 100 % | 17 | 100 % | 100 % | 49 |
| Average | 92.3 % | 100 % | 17 | 98.7 % | 99.7 % | 51 |
| SD | 6.7 % | 0 % | 5 | 1.2 % | 0.6 % | 9 |

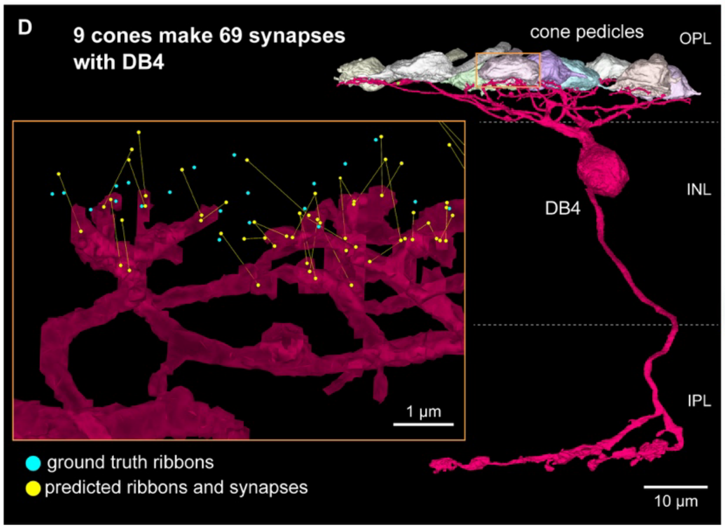


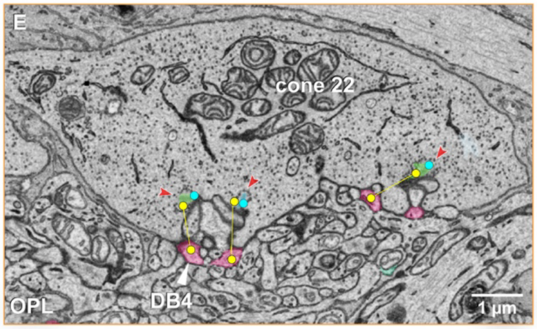


**F**

| Bipolars | synapse precision | synapse sensitivity | total (#) |
|---|---|---|---|
| DB4 | 87 % | 95 % | 73 |
| DB2 | 89 % | 89 % | 53 |
| Average | 88.0 % | 92.0 % | 63 |
| SD | 1.4 % | 4.2 % | 14 |

**Fig. S2. Further evaluation of excitatory and conventional synaptic detection. A.** Ribbon identification and synaptic prediction between cone 22 and FMB and IMB. 3D view shows cone 22 pedicle (white) and inner-ON (IMB, light orange) and outer-OFF (FMB, light green) bipolar cells. Cone 22 made 24 and 36 contacts with the IMB and FMB cells, respectively (one false positive ribbon was predicted for cone 22). Inset shows the dendritic terminals of the two bipolar cells. Ground truth identification of cone ribbons (blue dots) and predicted ribbons and synaptic connections (yellow dots and lines) are shown. **B**. Single EM layer view through cone 22 shows ribbon segmentation (ribbons in varied colors) and two predicted synaptic connections (yellow dots and lines) made with both IMB and FMB dendrites. **C**. Table shows precision and sensitivity for all midget bipolar connections made by cones 21, 22, and 58. **D**. As in **A**, but for a single diffuse bipolar cell (classified as a DB4 type) that was postsynaptic to cone 22 and eight additional cones. This DB4 cell made 69 contacts (both basal and invaginating contacts) with the nine cone pedicles. Inset conventions as in **A** show synaptic predictions for the connection with cone 22. **E**. Conventions as in **B**, showing three predicted basal contacts between cone 22 and this DB4 cell. **F.** Table shows precision and sensitivity of synaptic predictions for the DB4 cell and a DB2 cell that was postsynaptic to eight cones, including cone 22.

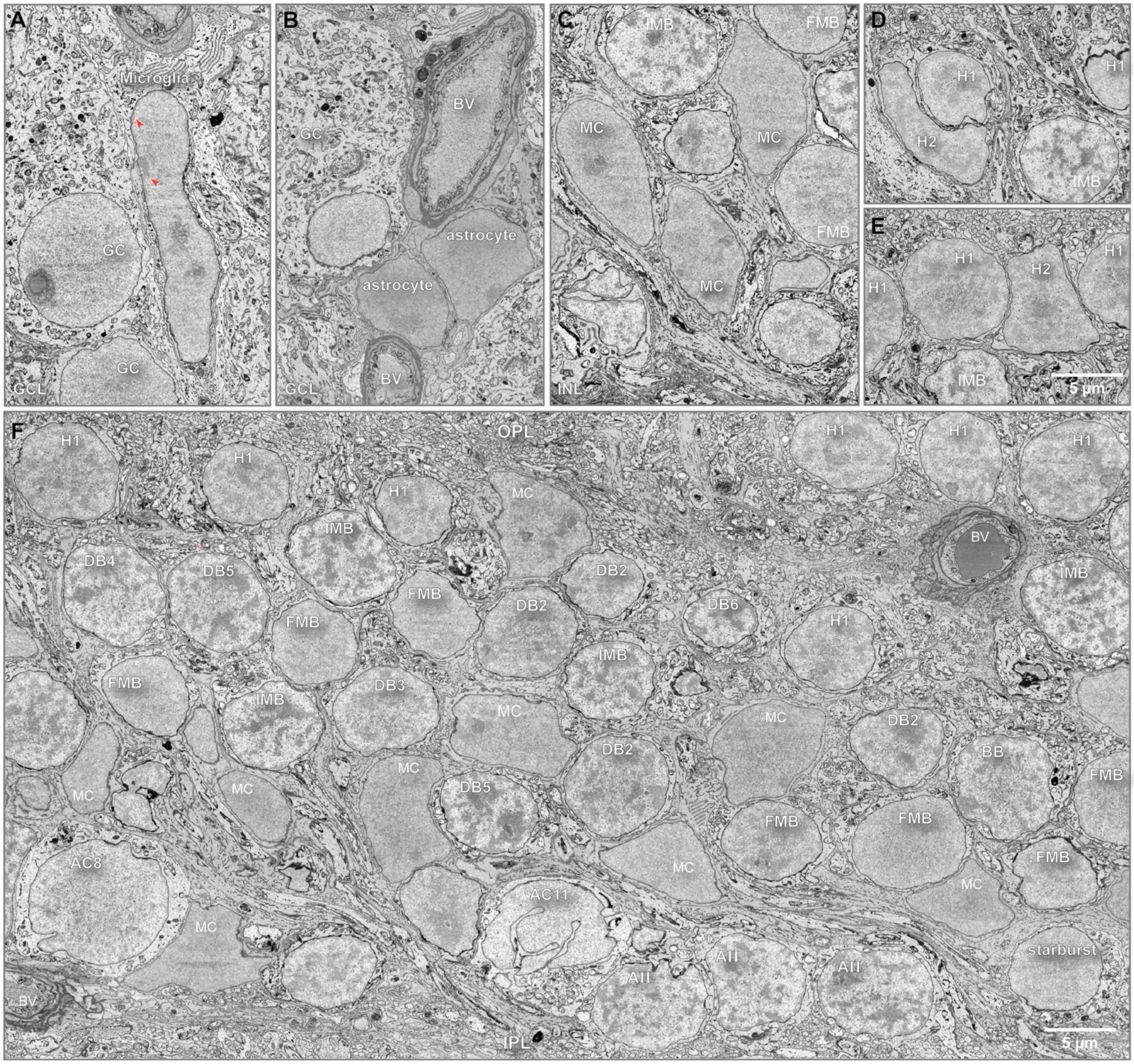


**Fig. S3. Distinguishing cell types by morphology of the nucleus.** Examples from single-layer EM view of nuclear morphology in identified (segmented) cell types in the inner nuclear and ganglion cell layers (INL, GCL). **A.** Microglial cells were easily recognized by their distinctly elongated nucleus, with heterochromatin accumulated at the nuclear margins (red arrowheads). All ganglion cells (GC) have round, euchromatic nuclei. **B.** Astrocytes were present in the GCL, and their cell bodies tended to accumulate around blood vessels; nuclei were relatively darkly stained and euchromatic. **C.** Müller glia cells (MC) were recognized by very large, ovoid or irregularly-shaped nuclei. Müller cell nuclei were euchromatic and moderately stained and similar in appearance to astrocyte nuclei but appeared larger. By contrast, the nuclei of bipolar cells (BC) are rounded. Invaginating-ON (IMB) midget bipolar cells show distinct heterochromatic nuclear staining marked by alternating very darkly and lightly stained regions. By contrast flat midget bipolar cells (FMB) show light euchromatic staining. **D** and **E.** H1 and H2 horizontal cells both show heterochromatic nuclear staining, but the dark-light contrast is less than that shown for the IMB cells. H1 cells have rounded nuclei, whereas the nuclei of H2 cells are irregularly shaped. **F.** Overview of nuclear patterning in the inner nuclear layer (INL). Like IMB cells, other inner stratifying-ON bipolar types (DB4, DB5, DB6 and BB) all show darkly stained, dense heterochromatic nuclear staining. However, unlike FMB cells the other outer stratifying OFF-bipolar cells (DB1, DB2, DB3) show lighter stained heterochromatic nuclei (a DB1 nucleus does not appear in this image), similar to H1 horizontal cells. Along the inner border of the INL amacrine cells display diverse nuclear morphologies. Starburst amacrine cells (see Fig. 5C) show round, lightly stained euchromatic nuclei; the starburst shown is an OFF-starburst with soma in the INL; ON-starbursts in the GCL show the same nuclear morphology as OFF-starburst cells. AII amacrine cells (Fig. 5A) are located at the border with the inner plexiform layer (IPL*)*. They show some scalloping of the nuclear envelope and are strongly heterochromatic, similar in appearance to ON bipolar cells. Other amacrine cell types are further distinguished by the degree of infolding of the nuclear envelope (crenulation) together with the distribution of chromatin. The small-field A11 amacrine cells (see Fig. 5E) show extremely infolded nuclear envelopes and light heterochromatic staining. By contrast with the AC11 cell, the large-field AC8 amacrine cell shows less crenulation and euchromatic nuclear staining. See *SI Appendix*, Figs. S6 and S8 for additional examples of distinctive nuclear morphology in amacrine cell types. BV, blood vessels.

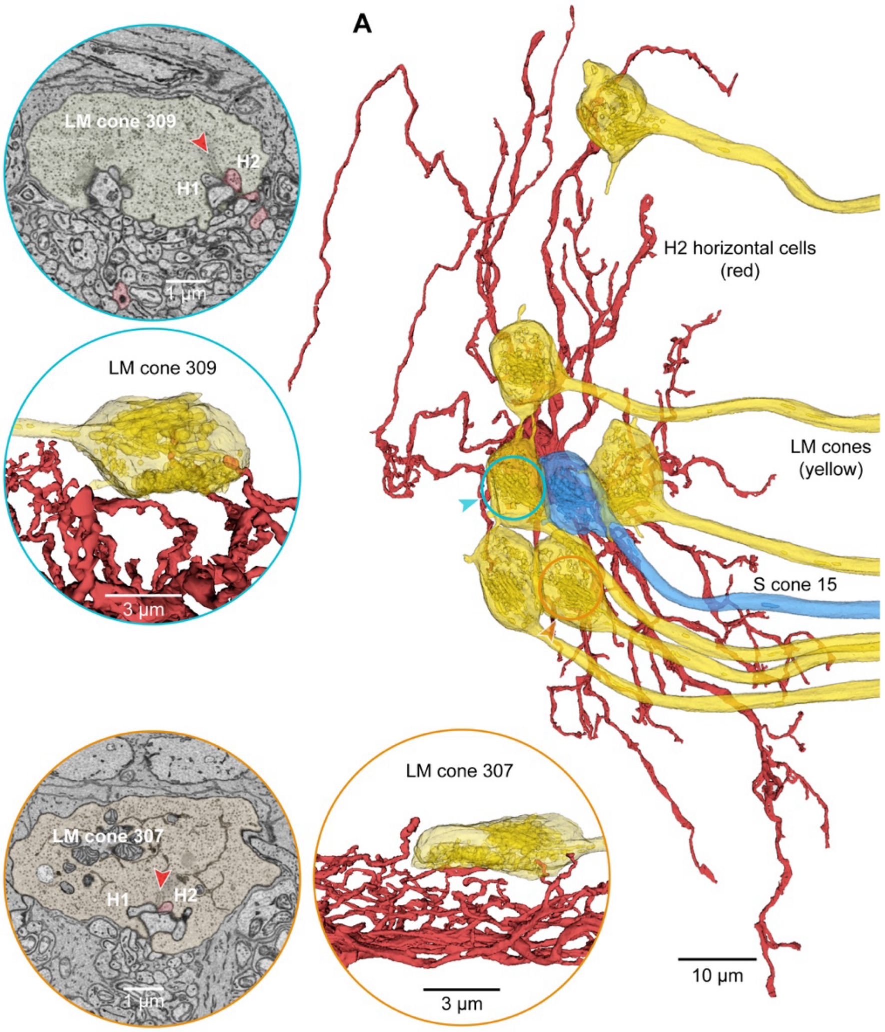


**Fig. S4. H2 horizontal cells only sparsely contact LM cones. A.** In HFseg1, H2 cells are sparsely branching and show little overlap with neighboring cells of the same type. A single H2 cell (red) is shown in relation to six LM cone pedicles (yellow) and one S cone (blue, S cone 15; see main Fig. 3 for S cone connectivity). The cone pedicles are shown in partial transparency. The blue-circled LM cone (LM cone 309; blue arrowhead) is shown in the insets at the upper left. This cone is contacted by only 1 lateral element from this H2 cell; the top inset shows a single-layer EM view, and the lower inset shows a 3D view. By contrast, H1 cells made 51 contacts with this cone, accounting for 98.1% of the lateral elements. A second orange-circled cone (LM cone 307; orange arrowhead) showed a similar pattern of horizontal cell connectivity, again making a single lateral contact with this H2 cell (1.8% of lateral elements), and 55 contacts with H1 cells (98.2% of lateral elements). This connection pattern was proofread for accuracy. Together with main Figure 3 the data analyzed thus far show that H1 cells receive major input from all three cone types.

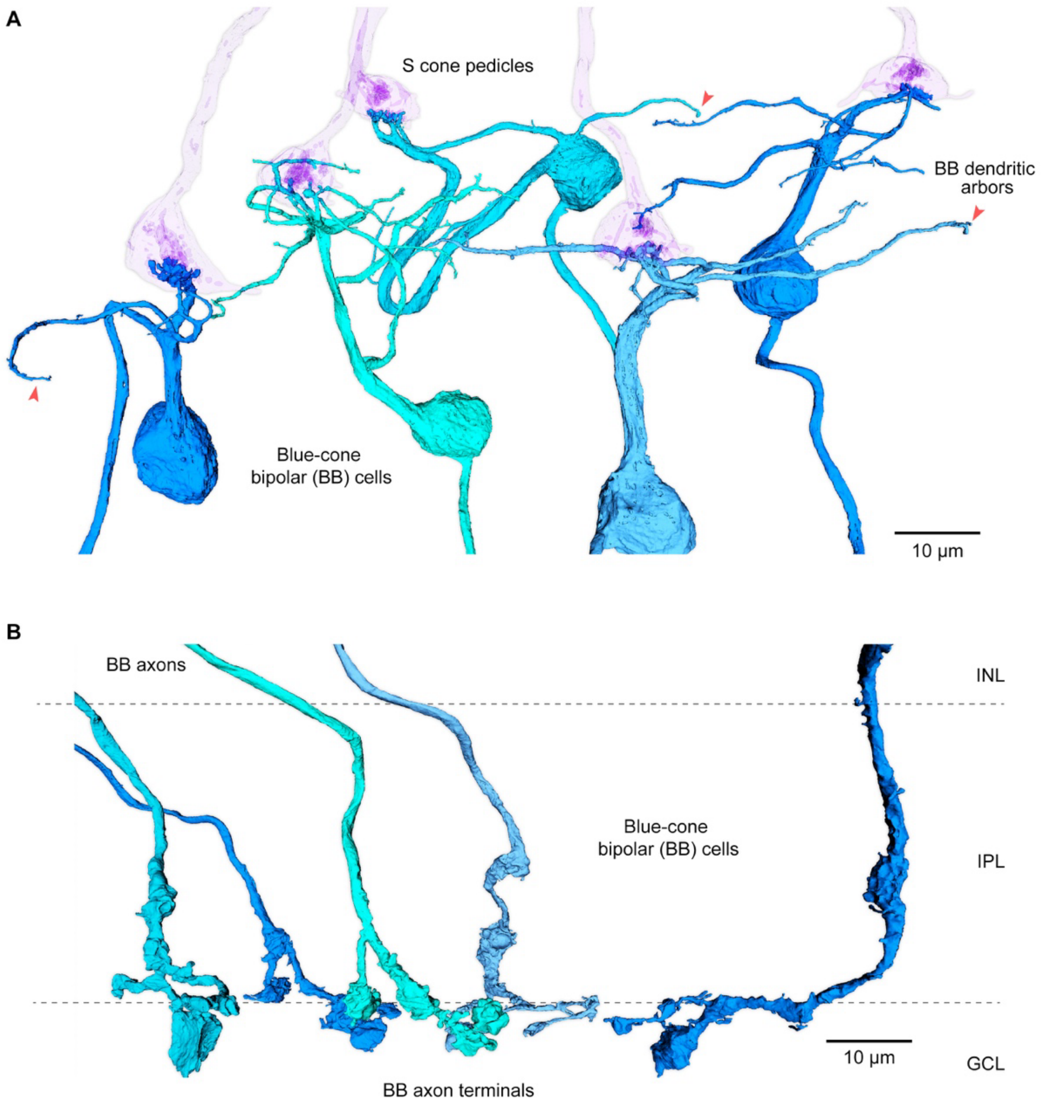


**Fig. S5. Blue cone bipolar cells (BB cells) show a distinctive dendritic morphology and axonal stratification. A**. Dendrites of BB cells primarily target S cone pedicles where they make invaginating contacts. Five S cones are shown in partial transparency (violet), tilted slightly off the horizontal plane. These S cones are contacted by the dendrites of five BB cells (in shades of blue); there is a tendency at this foveal location for each S cone to be presynaptic primarily to a single BB cell. Primary dendrites of the five BB cells are indicated by red arrowheads. **B.** The axon terminals of these five BB cells are shown in a vertical view (image is also rotated relative to the view in **A**). The axons terminate along the inner border of the IPL, often extending terminal varicosities across the outer border of the ganglion cell layer (GCL).

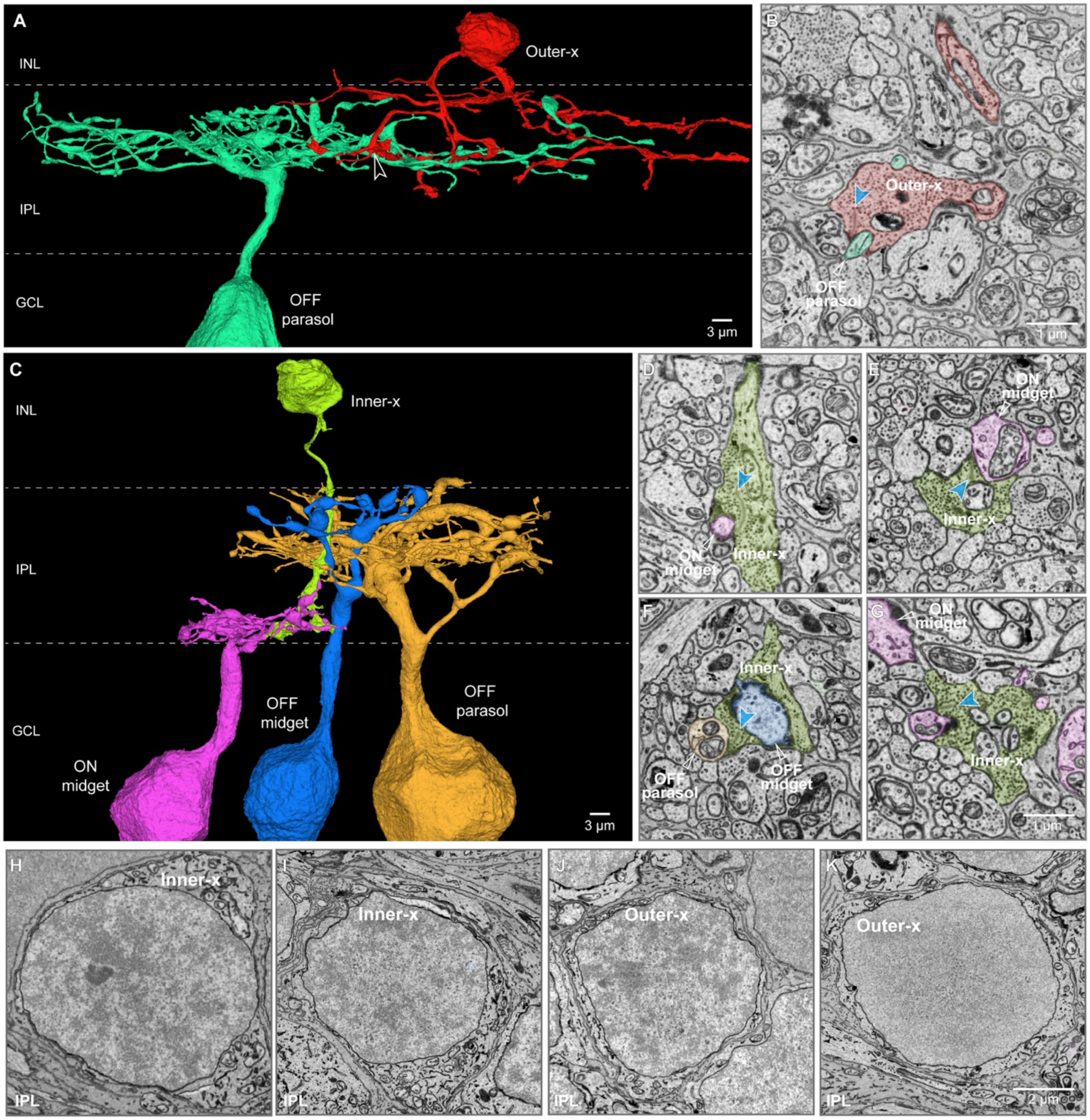


**Fig. S6. Inner- and Outer-x cells show amacrine like morphology, but, like bipolar cells make ribbon synapses in the IPL with ganglion cells.** The x cells vary in morphology and depth of stratification in the IPL and do not form any consistent spatial mosaic patterns typical of either bipolar or amacrine cells. **A**. An Outer-x cell with sparse broadly stratifying branches in the outer half of the IPL. It is presynaptic to an OFF-parasol GC (teal) (as well as other amacrine and ganglion cell processes, including OFF midget GCs (not shown)). **B**. Ribbon synapse (blue arrowhead) from the OFF-x cell shown in a to an OFF-parasol cell dendrite (indicated by open white arrowhead in a). **C**. Inner-x cell with a small axonal arbor stratified in the inner half of the IPL. **D-G**. Examples of ribbon synapses (blue arrowheads) made by the Inner-x cell shown in c with OFF parasol, OFF midget and ON midget GCs. **H-I**. Inner-x cells show light heterochromatic nuclear staining. **J-K**. Outer x-cells show both heterochromatic staining (**J**), like that of outer DB cells, and light euchromatic staining (**K**), like that of FMB cells, suggesting that the x cells may reflect modifications in the morphology of some number of bipolar types including IMB, FMB, and DB cells.

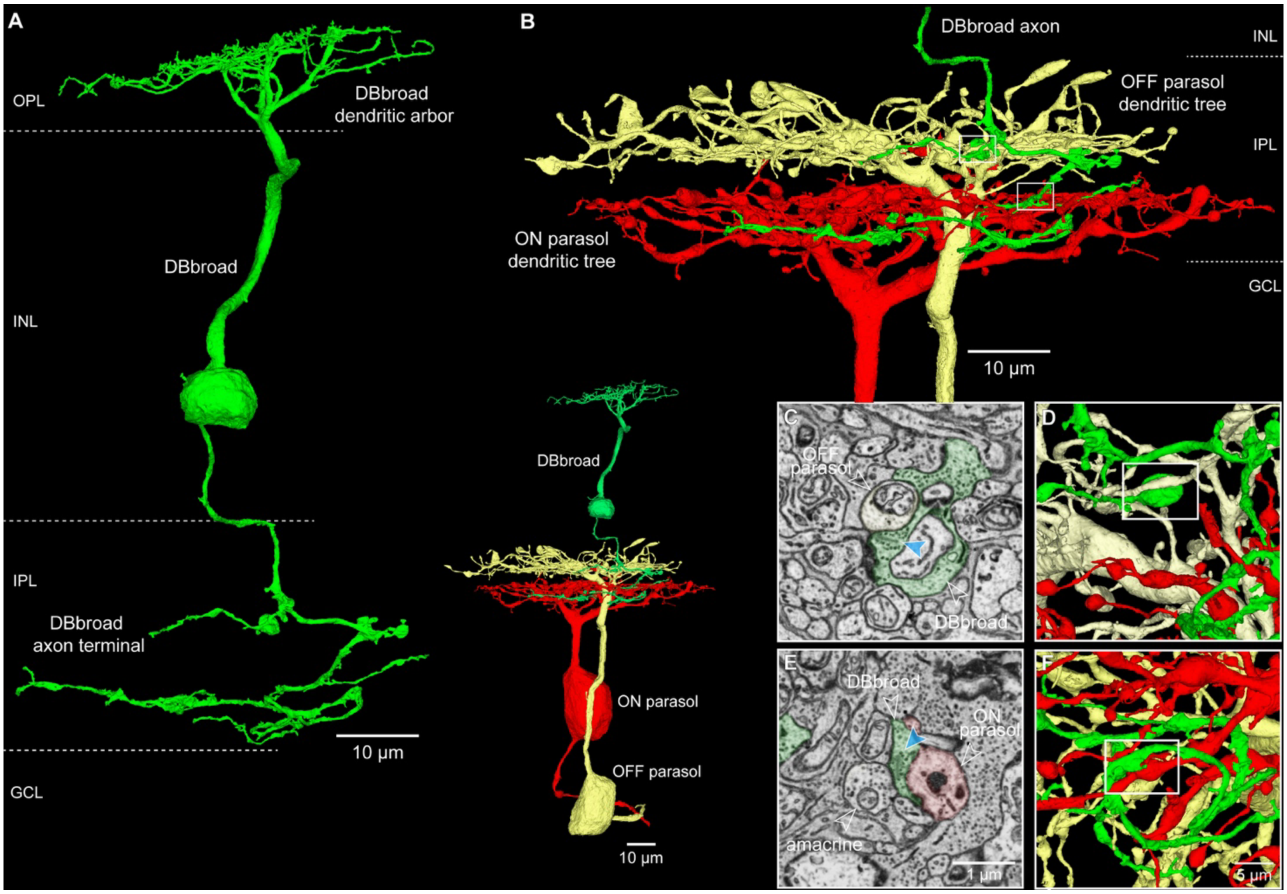


**Fig. S7. Distinctive morphology of DBbroad cells.** DBbroad cells do not appear to correspond to any previously identified cone bipolar cell type in primate retina. These cells have broadly stratifying axons and make sparse ribbon synapses in both the outer (OFF) and inner (ON) portions of the inner plexiform layer. **A.** The axonal arbor of the DBbroad cell is large and sparsely branching and does not show distinctive or large synaptic varicosities. **B**. This DBbroad cell is presynaptic to both an outer-OFF and an inner-ON parasol ganglion cell; a vertical view of the three cells is shown in the inset at center. At the upper right is a zoomed view of the axonal arbor of the DBbroad cell and the dendritic trees of the two parasol cells (ON cell, red; OFF cell, yellow). **C-D**. Ribbon synapse (blue arrowhead) made with the OFF parasol cell (yellow; open arrowhead) shown in a single section view. Location of this synapse in 3D view is outlined by the white box in **D**. **E-F**. Synapse made with the ON parasol cell (red; open arrowhead), and an amacrine cell (open arrowhead) in **E** shown in a single section view. Location of this synapse in 3D view is outlined by the white box in **F**. Approximate synaptic locations shown in **D** and **F** are also indicated by the white boxes in **B**.

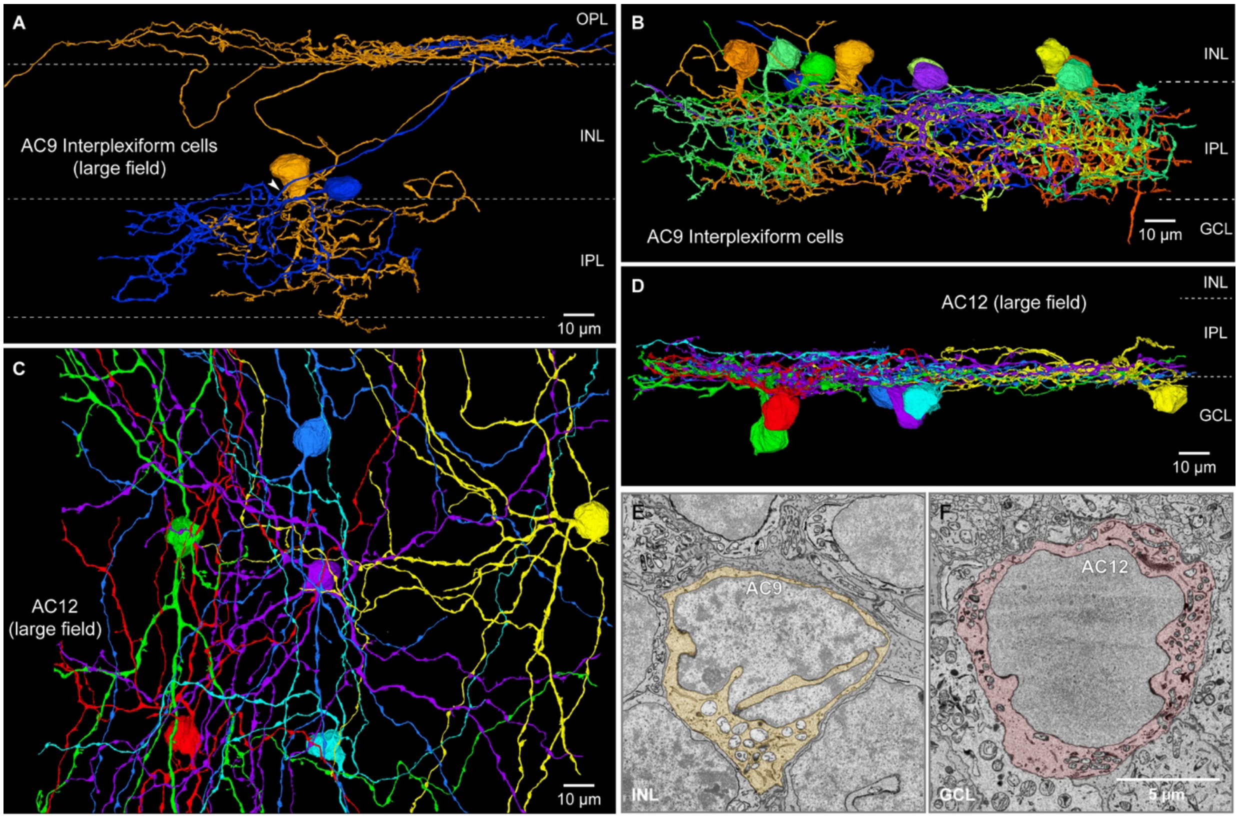


**Fig. S8. Additional amacrine cell populations in the fovea. A.** AC9 is a large-field amacrine type that shows the distinctive morphology of interplexiform cells (cells with processes in both outer and inner plexiform layers). A single axon-like (white arrowhead) process extends vertically from a primary dendrite in the IPL into the OPL where it branches and extends laterally; the inner processes of the cells are large, sparsely branching, and tortuous, extending across the full depth of the IPL. The synaptic connections of the outer and inner processes remain to be investigated. **B.** Vertical view of the dendritic trees of ten AC9 cells in the IPL (complete demonstration of all axon-like processes and their synaptic connections will require additional proofreading of each cell). **C.** AC12 is a large-field amacrine type with cell bodies displaced to the ganglion cell layer (GCL). Shown here a horizontal view of six A12 cells; the dendrites are thick, radiating, and moderately branched, similar in appearance to the AC8 cells shown in Fig. 5G. **D.** Vertical view of the same mosaic of AC12 cells; these cells stratify broadly over the inner third of the IPL where they costratify with and synapse onto the axon terminals of midget bipolar (IMB) cells (not shown). Details of the synaptic relationships of this cell type and its relationship to amacrine types previously identified in human and non-human primate retina remain to be clarified. AC9 and AC12, like other amacrine types show distinctive nuclear shapes and staining patterns. **E.** AC9 shows a highly infolded nuclear envelope combined with dark heterochromatic nuclear staining. **F.** AC12 shows a scalloped-irregular nuclear envelope combined with light euchromatic nuclear staining.

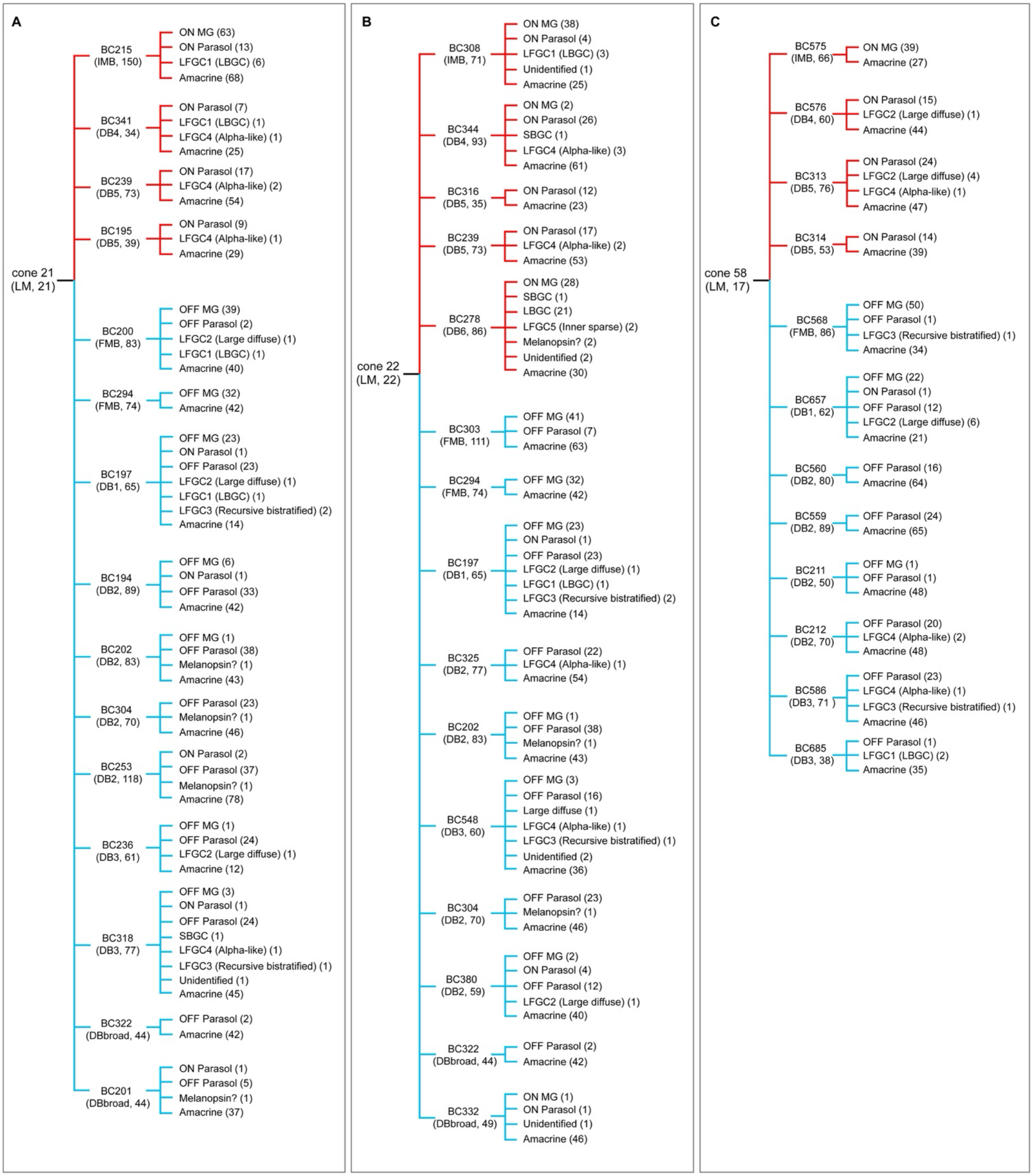


Note: as examples, cone 21 (LM, 21) indicates (cone type, # of total ribbon), BC215 (IMB, 150) indicates (bipolar type, # of total synapse).

**Fig. S9. Vertical excitatory connectome: outgoing synaptic contacts from all bipolar cells connected to 3 LM cones.** The bipolar type and number of synapses is shown in parentheses (e.g.,

BC 215 (IMB (cell type), 150 (outgoing synapses)). Synapses directed to ganglion cells were identified by type. Synapses directed to amacrine cells were summed and not yet divided by amacrine type). It can be noted that the total number of cells and synapses arising from OFF-bipolar types (blue connecting lines) is much higher than that for ON-bipolar cells (red-connecting lines); moreover, OFF bipolar cells are presynaptic to both ON and OFF ganglion cells whereas ON bipolar cells connect exclusively to inner ON ganglion cell types. It can also be noted that midget and parasol ganglion cells share significant convergent input from multiple bipolar cell types. These factors account for the high number of synapses in OFF visual pathways compared to ON pathways, as shown in Fig. 10C. Note that the three LM cones are shown in Fig. 2A (gold, white asterisks).

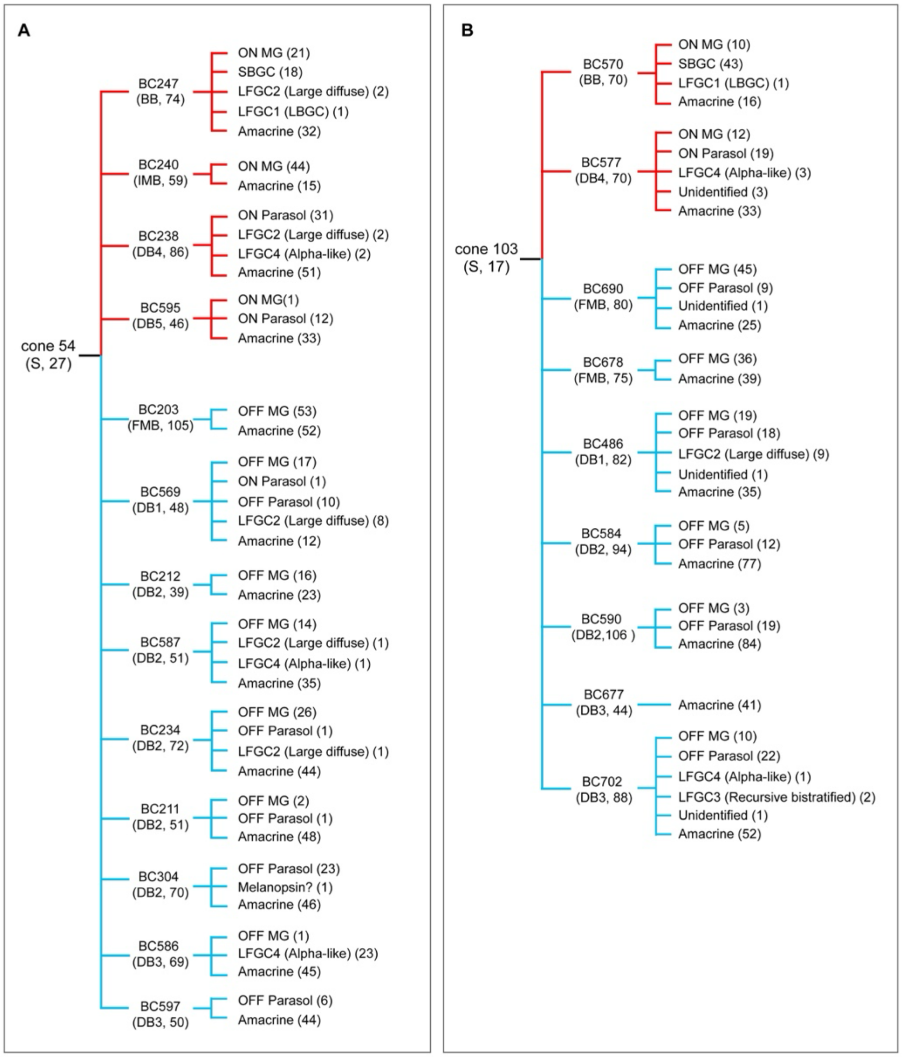


Note: as examples, cone 54 (S, 27) indicates (cone type, # of total ribbon), BC247 (BB, 74) indicates (bipolar type, # of total synapse).

**Fig. S10. Vertical excitatory connectome: outgoing synaptic contacts from all bipolar cells connected to the 2 S cones.** The bipolar type and number of synapses is shown in parentheses (e.g., BC 247 (BB (cell type), 74 (outgoing synapses)). Note that the two S cones are shown in Fig. 2A (blue, white asterisks).

**A** Summary

| Type | Total (n) | Total (%) |
|---|---|---|
| Photoreceptor | 337 | 11.2 % |
| Horizontal cell | 268 | 8.9 % |
| Bipolar cell | 912 | 30.3 % |
| Amacrine cell | 293 | 9.7 % |
| Ganglion cell | 599 | 19.9 % |
| Müller glia | 555 | 18.4 % |
| Astrocyte | 32 | 1.1 % |
| Microglia | 15 | 0.5 % |
| Total | 3012 | 100% |

**B** Non-neuronal cells

| Type | Total (n) | Total (%) |
|---|---|---|
| Müller glia | 555 | 92.2 % |
| Astrocytes | 32 | 5.3 % |
| Microglia | 15 | 2.5 % |
| Total | 602 | 100 % |

**C** Cone photoreceptors

| Type | Total (n) | Total (%) |
|---|---|---|
| LM cone | 296 | 94.6 % |
| S cone | 17 | 5.4 % |
| Total | 313 | 100% |

**D** Rod and cone photoreceptors

| Type | Total (n) | Total (%) |
|---|---|---|
| Cone | 313 | 92.9 % |
| Rod | 24 | 7.1 % |
| Total | 337 | 100% |

**E** Horizontal cells

| Type | Total (n) | Total (%) |
|---|---|---|
| H1 | 256 | 95.5 % |
| H2 | 12 | 4.5 % |
| Total | 268 | 100% |

**F** Bipolar cells (BCs)

| | Type | Total (n) | Total (%) |
|---|---|---|---|
| OFF | FMB | 302 | 33.1 % |
| | DB1 | 54 | 5.9 % |
| | DB2 | 92 | 10.1 % |
| | DB3 | 41 | 4.5 % |
| | Outer-x | 15 | 1.6 % |
| | DBbroad | 18 | 2.0 % |
| ON | IMB | 264 | 28.9 % |
| | DB4 | 34 | 3.7 % |
| | DB5 | 45 | 4.9 % |
| | DB6 | 12 | 1.3 % |
| | BB | 17 | 1.9 % |
| | RB | 6 | 0.7 % |
| | Inner-x | 5 | 0.5 % |
| | Unidentified | 7 | 0.8 % |
| | Total | 912 | 100% |

**G** Amacrine cells (ACs)

| Type | Total (n) | Total (%) | |
|---|---|---|---|
| AC2 (AII, small) | 75 | 25.6 % | 64.5 % |
| AC3 (small) | 29 | 9.9 % | |
| AC6 (small) | 24 | 8.2 % | |
| AC11 (small) | 22 | 7.5 % | |
| AC5 (small) | 20 | 6.8 % | |
| AC4 (small) | 19 | 6.5 % | |
| AC7 (SAC, large) | 17 | 5.8 % | 35.5 % |
| AC16 (large) | 11 | 3.8 % | |
| AC9 (Interplexiform, large) | 11 | 3.8 % | |
| AC10 (large) | 10 | 3.4 % | |
| AC8 (large) | 10 | 3.4 % | |
| AC19 (large) | 8 | 2.7 % | |
| AC1 (A1?, large) | 6 | 2.0 % | |
| AC12 (large) | 6 | 2.0 % | |
| AC15 (large) | 6 | 2.0 % | |
| AC17 (large) | 6 | 2.0 % | |
| AC13 (large) | 5 | 1.7 % | |
| AC18 (large) | 5 | 1.7 % | |
| AC14 (large) | 3 | 1.0 % | |
| Total | 293 | 100 % | |

**H** Ganglion cells (GCs)

| | Type | Total (n) | Total (%) | |
|---|---|---|---|---|
| Major GCs | OFF midget | 280 | 46.7 % | 95.8 % |
| | OFF parasol | 13 | 2.2 % | |
| | ON midget | 256 | 42.7 % | |
| | ON parasol | 13 | 2.2 % | |
| | SBGC | 12 | 2.0 % | |
| Large Field GCs | LFGC1 (LBGC) | 6 | 1.0 % | 4.2 % |
| | LFGC2 (Large diffuse) | 6 | 1.0 % | |
| | LFGC3 (Recursive bistratified) | 4 | 0.7 % | |
| | LFGC4 (Inner/outer smooth monostratified) | 5 | 0.8 % | |
| | LFGC5 (Inner sparse) | 4 | 0.7 % | |
| | Total | 599 | 100 % | |

**Table S1. Identification of cell populations.**

**a**

**Cone 21 (LM cone 21)**

| | NO | GC types | Total synapse (#) | Total percent (%) | |
|---|---|---|---|---|---|
| Major GCs | 1 | ON MG | 63 | 13.8 % | 94.5 % |
| | 2 | OFF MG | 105 | 23.0 % | |
| | 3 | ON Parasol | 51 | 11.4 % | |
| | 4 | OFF Parasol | 213 | 46.1 % | |
| | 5 | SBGC | 1 | 0.2 % | |
| Large Field GCs | 6 | LFGC1 (LBGC) | 9 | 2.0 % | 5.5 % |
| | 7 | LFGC2 | 3 | 0.7 % | |
| | 8 | LFGC3 | 3 | 0.7 % | |
| | 9 | LFGC4 | 5 | 1.1 % | |
| | 10 | LFGC5 | 0 | 0.0 % | |
| | 11 | Melanopsin? | 4 | 0.9 % | |
| | 12 | Unidentified | 1 | 0.2 % | |
| | | Total | 457 | 100% | |

**b**

**Cone 22 (LM cone 22)**

| | NO | GC types | Total synapse (#) | Total percent (%) | |
|---|---|---|---|---|---|
| Major GCs | 1 | ON MG | 69 | 16.0 % | 88.4% |
| | 2 | OFF MG | 101 | 23.7 % | |
| | 3 | ON Parasol | 65 | 15.1 % | |
| | 4 | OFF Parasol | 139 | 33.1 % | |
| | 5 | SBGC | 2 | 0.5 % | |
| Large Field GCs | 6 | LFGC1 (LBGC) | 25 | 5.8 % | 11.6 % |
| | 7 | LFGC2 | 3 | 0.7 % | |
| | 8 | LFGC3 | 3 | 0.7 % | |
| | 9 | LFGC4 | 7 | 1.6 % | |
| | 10 | LFGC5 | 2 | 0.5 % | |
| | 11 | Melanopsin? | 4 | 0.9 % | |
| | 12 | Unidentified | 6 | 1.4 % | |
| | | Total | 431 | 100% | |

**c**

**Cone 58 (LM cone 58)**

| | NO | GC types | Total synapse (#) | Total percent (%) | |
|---|---|---|---|---|---|
| Major GCs | 1 | ON MG | 39 | 13.8 % | 93.3 % |
| | 2 | OFF MG | 73 | 25.8 % | |
| | 3 | ON Parasol | 54 | 19.1 % | |
| | 4 | OFF Parasol | 98 | 34.6 % | |
| | 5 | SBGC | 0 | 0.0 % | |
| Large Field GCs | 6 | LFGC1 (LBGC) | 2 | 0.7 % | 6.7 % |
| | 7 | LFGC2 | 11 | 3.9 % | |
| | 8 | LFGC3 | 2 | 0.7 % | |
| | 9 | LFGC4 | 4 | 1.4 % | |
| | 10 | LFGC5 | 0 | 0.0 % | |
| | 11 | Melanopsin | 0 | 0.0 % | |
| | 12 | Unidentified | 0 | 0.0 % | |
| | | Total | 283 | 100% | |

**d**

**Cone 54 (S cone 6)**

| | NO | GC types | Total synapse (#) | Total percent (%) | |
|---|---|---|---|---|---|
| Major GCs | 1 | ON MG | 66 | 19.4 % | 94.4 % |
| | 2 | OFF MG | 129 | 37.9 % | |
| | 3 | ON Parasol | 44 | 12.9 % | |
| | 4 | OFF Parasol | 64 | 18.8 % | |
| | 5 | SBGC | 18 | 5.4 % | |
| Large Field GCs | 6 | LFGC1 (LBGC) | 1 | 0.3 % | 5.6 % |
| | 7 | LFGC2 | 14 | 4.1 % | |
| | 8 | LFGC3 | 0 | 0.0 % | |
| | 9 | LFGC4 | 3 | 0.9 % | |
| | 10 | LFGC5 | 0 | 0.0 % | |
| | 11 | Melanopsin | 1 | 0.3 % | |
| | 12 | Unidentified | 0 | 0.0 % | |
| | | Total | 340 | 100% | |

**e**

**Cone 103 (S cone 5)**

| | NO | GC types | Total synapse (#) | Total percent (%) | |
|---|---|---|---|---|---|
| Major GCs | 1 | ON MG | 22 | 7.2 % | 92.8 % |
| | 2 | OFF MG | 118 | 38.8 % | |
| | 3 | ON Parasol | 19 | 6.3 % | |
| | 4 | OFF Parasol | 80 | 26.3 % | |
| | 5 | SBGC | 43 | 14.1 % | |
| Large Field GCs | 6 | LFGC1 (LBGC) | 1 | 0.3 % | 7.2 % |
| | 7 | LFGC2 | 9 | 3.0 % | |
| | 8 | LFGC3 | 2 | 0.7 % | |
| | 9 | LFGC4 | 4 | 1.3 % | |
| | 10 | LFGC5 | 0 | 0.0 % | |
| | 11 | Melanopsin | 0 | 0.0 % | |
| | 12 | Unidentified | 6 | 2.0 % | |
| | | Total | 304 | 100% | |

**Table S2. Total excitatory synaptic output from a single cone. Links to cells in NeuroMaps.** The second columns contain clickable links to view the various neurons that make up the HFseg1 volume. Note that two S cones and three LM cones are demonstrated in Fig. 2A.

| | |
|---|---|
| Non-neuronal cells<br>(related to *SI Appendix*, Fig. S1) | Microglia cells<br>Astrocytes<br>Müller Glia cells |
| Horizontal cells<br>(related to Fig. 3A, 3B) | H1 Horizontal cells<br>H2 Horizontal cells |
| Photoreceptors<br>(related to Fig. 2) | LM/S Cones + Rods |
| Bipolar cells<br>(related to Fig. 4) | IMB (ON midget bipolar cells)<br>FMB (OFF midget bipolar cells)<br>Blue cone bipolar cells (BB cells)<br>ON diffuse bipolar cells (DB4, DB5, DB6)<br>OFF diffuse bipolar cells (DB1, DB2, DB3)<br>RB (rod bipolar)<br>Outer-x<br>Inner-x<br>Giant<br>DBbroad (diffuse bipolar type) |
| Amacrine cells<br>(related to Figs. 5, 9) | AC2 (AII amacrine cells)<br>AC7 (starburst amacrine cells)<br>AC 8<br>AC 11 |
| Ganglion cells<br>(related to Fig. 6) | ON and OFF midget ganglion cells and the midget circuit<br>ON/OFF parasol ganglion cells<br>Small bistratified ganglion cells<br>LFGC1 (Large bistratified)<br>LFGC2 (Large diffuse)<br>LFGC3 (Recursive bistratified)<br>LFGC4 (Smooth monostratified) |

**Table S3. Links to cells in NeuroMaps.** The second columns contain clickable links to view the various neuron and glial cell populations that comprise the HFseg1 volume.

**A**

| Model parameters | Value |
|---|---|
| Specific membrane resistance ($R_m$) | 12,000 Ohm−cm$^2$ |
| Axon resistivity ($R_i$) | 80 Ohm-cm |
| Cone soma diameter | 2.5 µm |
| Cone axon diameter | 1.6 µm |
| Cone axon length | 300 µm |
| Cone axon terminal diameter | 5 µm |
| Rod soma diameter | 2.5 µm |
| Rod axon diameter | 0.45 µm |
| Rod axon terminal diameter | 3 µm |
| Axon $V_{rev}$ | -60 mV |
| Compartment size | 0.05 lambda |
| Cone axon compartments | 9 |
| Rod axon compartments | 11 |
| Cone OS conductance | 880 pS |
| Background level | 10,000 Ph/µm$^2$/s |
| Stimulus level | 100,000 Ph/µm$^2$/s (monochromatic) |
| Stimulus duration | 10 ms flashes |
| Stimulus size | 1000 µm (diameter spot) |

**B**

| From data | Value |
|---|---|
| Total # of cones | 310 |
| Total # of rods | 23 |
| Average contact area from connectome | 0.55 µm |
| Connexin conductance | 15 pS |
| Assumed conductance | 75 pS (5 connexins) per average contact |
| Contact conductance | 136 pS/µm$^2$ |
| Average # of contacts per cone | 3.6 |
| Average LM pair coupling conductance | 115.9 pS |
| Average S-LM pair coupling conductance | 123.4 pS |

**C**

| From simulations | Value |
|---|---|
| Target L/M ratio | 1.7 |
| Actual L/M ratio | 1.697 (random assignment) |
| Cone terminal input resistance | ~0.4 Gohm |
| Cone $V_{rest}$ | ~ -40 mV (-40 +/- 1mV) |
| Measured average coupling conductance L/M pairs | 133 pS (n = 25) |
| Measured average coupling conductance S-LM pairs | 166 pS (n = 15) |

**Table S4. Model parameters for cone-cone coupling simulation.** Model parameters used in Fig. 8.
